# Structure and inhibition of the sperm TMEM95–FIMP complex in mammalian fertilization

**DOI:** 10.64898/2026.05.14.724122

**Authors:** Pulan Liu, Rithik E. Castelino, Taylor R. Gierke, Anna J. Wood, Yonggang Lu, Shaogeng Tang

## Abstract

TMEM95 is a sperm acrosomal membrane protein essential for mammalian fertilization. How TMEM95 facilitates sperm–egg interaction has largely remained unknown. Analogous sperm fertilization proteins function as complexes, leading us to hypothesize that TMEM95 may have a binding partner on sperm. Here, we surveyed interactions between TMEM95 and individual proteins in a curated library of testis-expressed proteins using AlphaFold3. We identify FIMP, a fertilization-essential acrosomal membrane protein, as a high-confidence interaction partner of TMEM95. These two proteins form a high-affinity complex through their ectodomains. Using single-particle cryo-EM, we determine the structure of the human TMEM95–FIMP ectodomain complex at high resolution. An aromatic motif of FIMP binds to a conserved surface of TMEM95, and amino acid substitutions within this motif ablate the TMEM95-binding activity of FIMP. We isolate an anti-TMEM95 antibody, termed 3A02, that binds to human and murine TMEM95 and disrupts the interaction between TMEM95 and FIMP. By determining the cryo-EM structure of human TMEM95 bound to the 3A02 fragment antigen-binding region, we find that 3A02 recognizes the FIMP-binding site on TMEM95. 3A02 inhibits fusion of murine sperm with eggs, independent of antibody size, suggesting that the TMEM95–FIMP interface is critical for sperm–egg interaction. Together, these results establish the human sperm TMEM95–FIMP complex and suggest that a FIMP-mediated interaction of TMEM95 facilitates membrane fusion during mammalian fertilization.

## Introduction

Fertilization is a central event of sexual reproduction. Several sperm proteins, including IZUMO1, SPACA6, TMEM81, FIMP, TMEM95, DCST1, DCST2, and SOF1, have been identified as essential factors in mammalian sperm–egg interactions ^1–12^. Among them, IZUMO1, SPACA6, and TMEM81 assemble into a heterotrimeric complex that interacts with the egg receptor, JUNO ^9,13,14^. DCST1 and DCST2 are suggested to form a heterodimeric complex to mediate fertilization ^5,8^. These recent studies suggest that the fertilization-essential sperm proteins work in concert to drive sperm–egg interaction. TMEM95 (transmembrane protein 95) facilitates membrane fusion during fertilization through an interaction with a putative egg receptor. The TMEM95 ectodomain constitutes a large, conserved surface with a solvent-accessible surface area comparable to those of protein interfaces that mediate protein–protein interactions ^7^. TMEM95 shares homology with IZUMO1 and SPACA6 ^15^, two proteins that co-assemble with TMEM81 on sperm. This structural evidence led us to hypothesize that, in addition to its egg-binding activity, TMEM95 also engages a sperm-expressed binding partner.

In this study, we aimed to identify a binding partner of TMEM95 on sperm and determine its role in mammalian fertilization. A previous proteomic study did not detect interactions between TMEM95 and sperm proteins essential for fertilization ^16^. Here, using *in silico* screening, biochemical, and structural biology approaches, we found that FIMP (fertilization influencing membrane protein) forms a high-affinity complex with TMEM95. Prior work has demonstrated that FIMP is a testis-specific, single-pass acrosomal membrane protein that relocalizes to the equatorial segment of the sperm head, where sperm–egg membrane fusion occurs ^17^. *Fimp* knockout mice show severe subfertility and the knockout sperm, despite normal expression and localization of IZUMO1, can bind to but do not fuse with, eggs ^17^. Collectively, these observations pointed to a role of TMEM95 and FIMP in sperm–egg interaction.

Here, we identified a novel sperm acrosomal membrane complex of TMEM95 and FIMP that drives sperm-egg interactions, defining FIMP as the partner of TMEM95 essential for mammalian fertilization. We determined a cryo-EM structure of the human TMEM95–FIMP ectodomain complex together with the fragment antigen-binding (Fab) of two anti-TMEM95 antibodies, 3A01 and 6B08. In this structure, we found that a conserved aromatic motif of FIMP interacts with an evolutionarily conserved, positively charged surface of TMEM95 previously shown to be important for egg binding ^7^. We isolated a third anti-TMEM95 antibody, a near-germline antibody termed 3A02, which cross-reacts with human and murine TMEM95 orthologs. We determined the cryo-EM structure of the human TMEM95 ectodomain with the Fab fragments of 3A02 and 6B08, revealing that 3A02 binds directly to the FIMP-binding surface of TMEM95. We further showed that 3A02 disrupts TMEM95–FIMP complex formation and inhibits sperm–egg fusion during murine fertilization. Alongside the IZUMO1–SPACA6–TMEM81 and DCST1–DCST2 complexes, our study identified TMEM95–FIMP as a third, distinct sperm membrane protein complex and demonstrates that the FIMP-mediated interaction with TMEM95 plays an essential role in mammalian sperm–egg membrane fusion.

## Results

### TMEM95 and FIMP ectodomains form a high-affinity complex

Motivated by a hypothesis that TMEM95 interacts with a sperm-expressed binding partner, we interrogated protein–protein interactions between TMEM95 and individual proteins in a curated library of 4,532 human testis-enriched secreted and membrane proteins. We screened human TMEM95 ectodomain (Fig. 1a) against individual proteins from the library with AlphaFold3 (Fig. 1b), predicting all binary complexes of TMEM95 and ranking them from high to low confidence based on minimum predicted alignment error (min-PAE) (Fig. 1c) and mean interface predicted template modeling-score (ipTM) scores (Extended Data Fig. 1b). The TMEM95–FIMP pair ranked first by min-PAE across all candidates and exhibited an ipTM score within the top 0.45% of predictions, consistent with a previous report ^14^. The other sperm membrane proteins essential for fertilization, IZUMO1, SPACA6, TMEM81, DCST1, DCST2, TMDD1 ^18^, and FAM187A ^18,19^, were not among the top hits (Fig. 1c and Extended Data Fig. 1a).

**Fig. 1.**
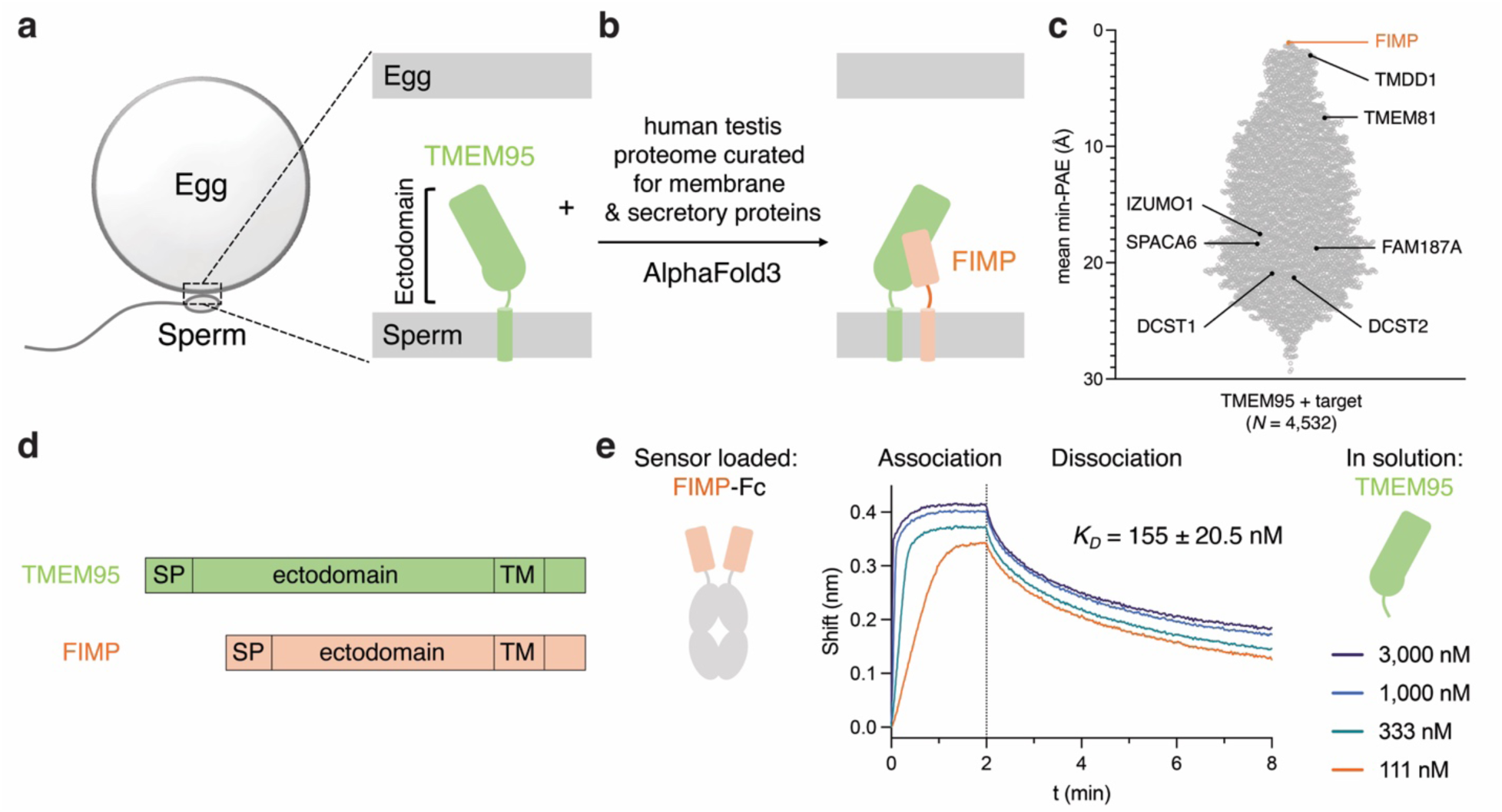
The human TMEM95 and FIMP ectodomains form a high-affinity complex. (a) Cartoon schematic showing sperm TMEM95 localized to the fertilization synapse. (b) AlphaFold3 screen of the TMEM95 ectodomain against a curated library of human testis-expressed, membrane-bound or secreted proteins. (c) Violin plot of mean minimum predicted aligned error (min-PAE) scores from 4,532 predicted protein–protein interactions. FIMP is the top-ranked candidate; additional sperm fertilization-essential proteins are labeled. (d) Domain organization of TMEM95 and FIMP: signal peptide (SP), ectodomain, and transmembrane domain (TM). (e) Biolayer interferometry traces showing binding of sensor-loaded FIMP-Fc to TMEM95 ectodomain at 3,000 nM, 1,000 nM, 333 nM, and 111 nM. Association, 2 min; dissociation, 6 min.

To examine the interaction of TMEM95 and FIMP *in vitro*, we recombinantly expressed and purified their ectodomains (Fig. 1d). Using biolayer interferometry (BLI), we found that a fusion protein of human FIMP ectodomain with human immunoglobulin G fragment crystallizable region (FIMP-Fc) binds to the human TMEM95 ectodomain in a concentration-dependent manner, with a *K_D_* of 155 ± 20.5 nM (Fig. 1e). Consistent with this, when murine TMEM95 is overexpressed on the surface of HEK293F cells, the murine FIMP-Fc (mFIMP-Fc) protein binds to the cell surface in a concentration-dependent manner, with an *EC_50_* of 140 ± 30 nM (Extended Data Fig. 2a,b). Together, these results indicate that TMEM95 and FIMP interact through their ectodomains at high affinity.

### FIMP interacts with the conserved surface of TMEM95

To understand the structural basis of the high-affinity interaction between TMEM95 and FIMP at high resolution, we analyzed this complex together with two previously characterized anti-TMEM95 monoclonal antibodies, 3A01 and 6B08 ^7^. Using BLI, we found that TMEM95 binds to FIMP-Fc, 3A01, and 6B08 simultaneously, indicating that they recognize three distinct, non-overlapping sites on TMEM95 (Fig. 2a). We therefore used the two antibodies to stabilize the TMEM95–FIMP complex and increase the particle size for cryo-EM structure determination.

**Fig. 2.**
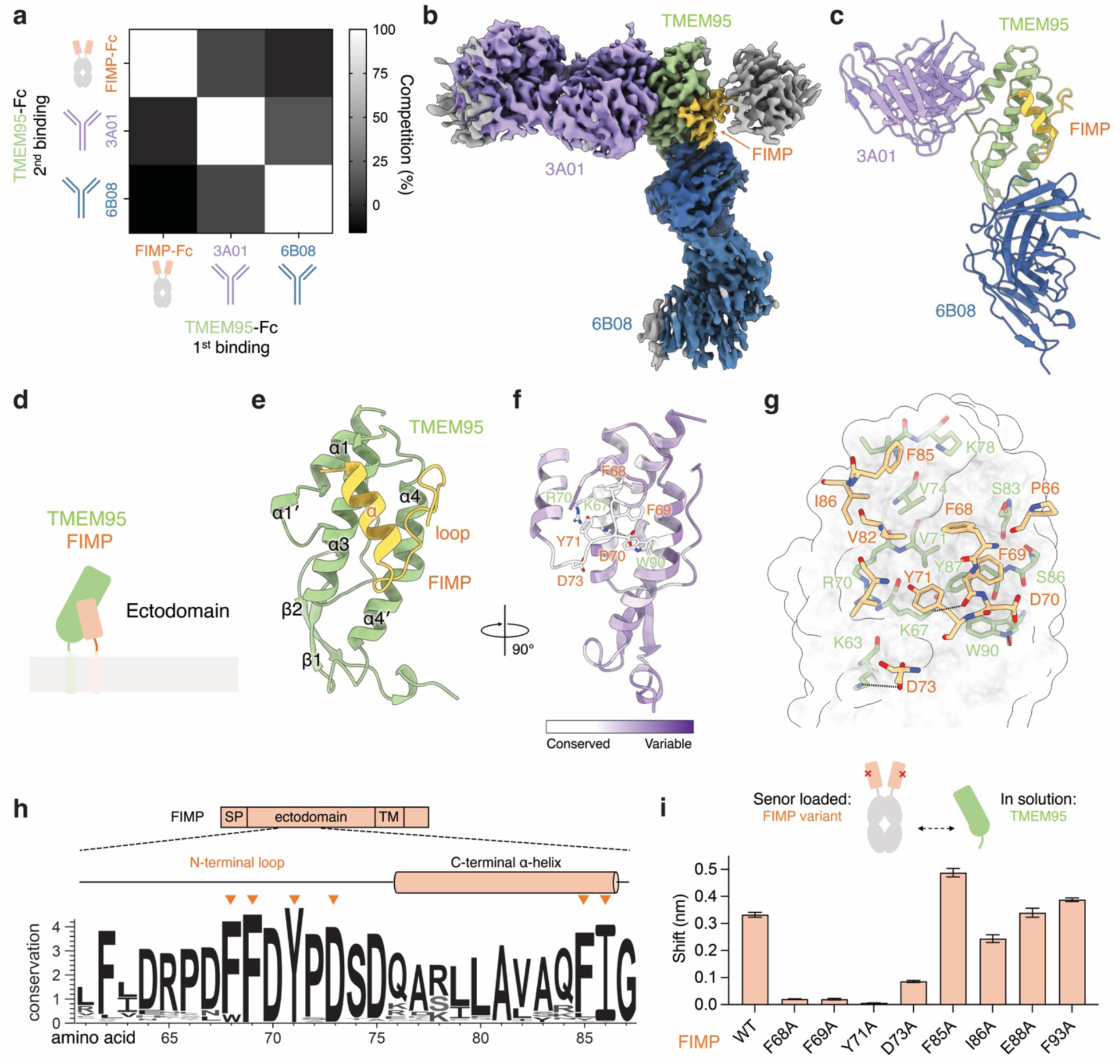
Structure of the human TMEM95–FIMP–3A01–6B08 complex, showing that FIMP interacts with the conserved surface of TMEM95. (a) Heat map summarizing competition between FIMP-Fc, 3A01 IgG, and 6B08 IgG for binding to sensor-immobilized TMEM95-Fc, showing that the three bind distinct regions of TMEM95. (b) Cryo-EM map of the human TMEM95–FIMP–3A01–6B08 complex. TMEM95, green; FIMP, yellow; 3A01, purple; 6B08, dark blue; and unmodeled Fc, grey. (c) Ribbon diagram of the cryo-EM structure of TMEM95–FIMP–3A01–6B08. (d) Cartoon schematic of the TMEM95–FIMP complex on the sperm membrane, highlighting the region of the ectodomains. (e) Ribbon diagram of the TMEM95–FIMP complex, in which the N-terminal loop and C-terminal α-helix of FIMP adopt a V-shaped conformation that interacts with the α3 and α4 helices of TMEM95. (f) Ribbon diagram colored by sequence conservation, with white indicating conserved regions and purple indicating variable regions. Side chains of key conserved interacting residues are shown. (g) Close-up view of the TMEM95–FIMP interaction interface with side chains of interacting residues shown. TMEM95, green with a semi-transparent white surface; FIMP, yellow. (h) WebLogo of FIMP orthologs showing sequence conservation across 213 BLAST-derived protein sequences. Letter height reflects amino acid frequency at each position, scaled by information content on the y-axis. (i) Biolayer interferometry shift values for sensor-loaded FIMP-Fc or its variants binding to TMEM95 at the 2-min association time point. Data are plotted as mean ± standard deviation from three replicate binding experiments.

A challenge we initially encountered was that the FIMP-Fc protein expresses poorly. To enhance protein expression, we generated a series of FIMP-Fc variants and found that single amino acid substitutions, F85A, I86A, or F93A, each drastically increased expression (Extended Data Fig. 2c). As these FIMP variants bind to TMEM95 comparably to wildtype (Extended Data Fig. 4a,b), we therefore used FIMP^F93A^ for subsequent structural determination. We designed an Fc-fusion protein in which one arm contains the TMEM95 ectodomain fused to Fc “knobs” and the other contains the FIMP^F93A^ ectodomain fused to Fc “holes”, with the knobs-into-holes interaction forming a covalent heterodimeric Fc (Extended Data Fig. 2e). We assembled a complex of TMEM95–FIMP^F93A^-Fc with 3A01 Fab and 6B08 Fab, purified to homogeneity (Extended Data Fig. 2d,e,f,g), and determined its structure by single-particle cryo-EM at 3.1 Å global resolution (Fig. 2b, Extended Data Fig. 3a,b,c,d,e,f, Table 1, and Extended Data Table 2). The resulting cryo-EM density map allowed assignment and model building of the TMEM95 and FIMP ectodomains, 3A01 fragments variable, and 6B08 fragments variable (Fig. 2c)

**Table 1.** Cryo-EM data collection, processing, and model validation statistics.

|  | Human TMEM95–FIMP<br>–3A01–6B08 | Human TMEM95<br>–3A02–6B08 |
| --- | --- | --- |
| **Data collection** |  |  |
| Voltage (kV) | 300 | 300 |
| Microscope | S2C2 TEMBeta<br>Titan Krios G3i | S2C2 TEMBeta<br>Titan Krios G3i |
| Camera | K3 (Gatan) | K3 (Gatan) |
| Magnification (calibrated) | 105,000 × | 105,000 × |
| Electron exposure (e<sup>−</sup>/Å<sup>2</sup>) | 50 | 50 |
| Exposure rate (e<sup>−</sup>/Å<sup>2</sup>/s) | 15 | 15 |
| Number of frames collected<br>per micrograph | 50 | 50 |
| Energy filter slit width | 20 eV | 20 eV |
| Automation software | EPU | EPU |
| Defocus range (μm) | -1.0 to -2.5 | -1.0 to -2.5 |
| Pixel size (Å) | 0.86 | 0.86 |
| **Overall map processing** |  |  |
| EMDB ID | EMD-77083 | EMD-77090 |
| Micrographs used | 6,856 | 7,238 |
| Symmetry imposed | C1 | C1 |
| Initial particle images | 4,048,830 | 1,991,260 |
| Final particle images | 538,876 | 115,272 |
| Resolution at 0.143 FSC of<br>masked reconstruction (Å) | 3.11 | 3.86 |
| Map sharpening B factor (Å<sup>2</sup>) | -123.7 | -155.2 |
| **Refinement** |  |  |
| PDB ID | 13IN | 13JC |
| Initial model used | Alphafold3 prediction | Alphafold3 prediction |
| Refinement package | Phenix v1.21 | Phenix v1.21 |
| **Map-model CC** |  |  |
| CC_mask | 0.88 | 0.84 |
| CC_box | 0.69 | 0.80 |
| CC_peaks | 0.63 | 0.70 |
| CC_volume | 0.86 | 0.82 |
| **Model composition** |  |  |
| Non-hydrogen atoms | 4,473 | 9124 |
| Protein residues | 594 | 1186 |
| **R.m.s. deviations** |  |  |
| Bond lengths (Å) | 0.002 | 0.003 |
| Bond angles (°) | 0.555 | 0.680 |
| B factors (Å<sup>2</sup>) |  |  |
| Protein | 62.86 | 92.25 |
| **Validation** |  |  |
| MolProbity score | 1.64 | 2.32 |
| Clashscore | 2.30 | 7.37 |
| Poor rotamers (%) | 2.54 | 3.89 |
| **Ramachandran plot** |  |  |
| Favored (%) | 95.35 | 92.65 |
| Allowed (%) | 4.48 | 7.27 |
| Disallowed (%) | 0.00 | 0.09 |
| Cβ outliers (%) | 0.00 | 0.00 |
| CaBLAM outliers (%) | 3.53 | 5.33 |

In this ternary complex, TMEM95 and FIMP ectodomains form a 1:1 stoichiometric complex, burying 563.8 Å^2^ of solvent-accessible surface area on TMEM95 (Fig. 2d,e and Extended Data Fig. 4d). TMEM95 adopts a fold similar to our previously reported apo structure ^7^, with an overall Cα root mean square deviation (r.m.s.d.) of 0.81 Å across 92 aligned residues (Extended Data Fig. 4c). The evolutionarily conserved, positively charged surface, formed by the α3 and α4 helices of TMEM95 undergoes a subtle conformational change to form a pocket that accommodates a conserved motif of FIMP (Fig. 2f,g and Extended Data Fig. 4c,d,e). In this interface, FIMP consists of a more conserved N-terminal loop (residues 62–75) followed by a short, less conserved C-terminal α-helix (residues 76–86), together adopting a V-shaped conformation that docks onto the conserved surface of TMEM95 (Fig. 2e,f,h and Extended Data Fig. 4d,e). Within the N-terminal loop of FIMP, F69, D70, and Y71 form hydrogen bonds with K67, W90, and R70 of TMEM95, respectively. This interaction is highly conserved across orthologs (Fig. 2f and Extended Data Fig. 4e). In the C-terminal α-helix of FIMP, V82, F85, and I86 engage in a hydrophobic interaction network with V71, V74, I77, and Y87 of TMEM95 (Fig. 2g and Extended Data Fig. 4e).

To examine whether this interface is critical for complex formation *in vitro*, we individually substituted amino acid residues of FIMP with alanine and evaluated binding to TMEM95 by BLI. We found that the F68A, F69A, and Y71A substitutions completely abolished complex formation, while substitutions outside the interface, E88A and F93A, had minimal effects (Fig. 2i and Extended Data Fig. 4a,b). These data provide strong evidence that the observed TMEM95–FIMP interface is required for their ectodomain heterodimerization.

### The anti-TMEM95 antibodies 3A01 and 6B08 recognize two distinct regions of TMEM95

We showed previously that the anti-TMEM95 antibodies 3A01 and 6B08 do not block binding of human sperm to hamster eggs but inhibit membrane fusion ^7^. In the ternary complex, we observed that 3A01 and 6B08 recognize TMEM95 outside the FIMP-binding surface, predominantly through their heavy-chain complementarity-determining regions (CDRs). 3A01 contacts the α1, α1′ helices, and the post-α1 loop of TMEM95 (Fig. 3a,b and Extended Data Fig. 3g), while 6B08 contacts the β-hairpin, the α4′ helix, and the post-α4′ loop (Fig. 3c,d and Extended Data Fig. 3h). Consistent with these assignments, substitution with alanine of R110 of TMEM95, which lies within the 6B08 epitope (Fig. 3e), reduces binding of 6B08 (Fig. 3d) but not 3A01 (Fig. 3e).

**Fig. 3.**
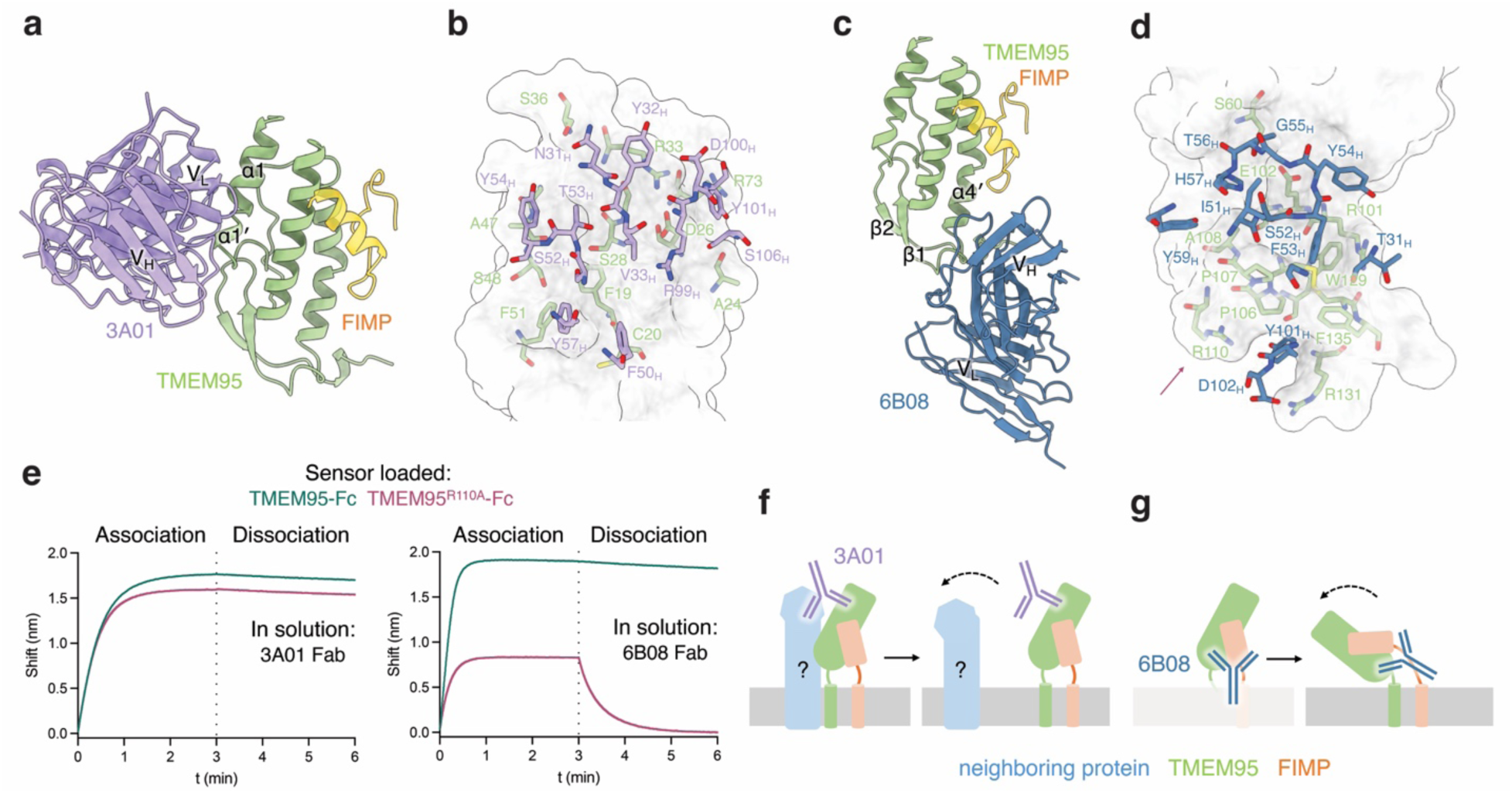
The anti-TMEM95 antibodies 3A01 and 6B08 recognize two distinct regions of TMEM95. (a) Ribbon diagram of 3A01–TMEM95–FIMP, in which the 3A01 variable regions interact with the α1 and α1′ helices of TMEM95. (b) Close-up view of the 3A01–TMEM95 interface. The interaction is mediated by hydrogen bonds and salt bridges between residues of the 3A01 heavy chain (V_H_: N31, V33, S52, T53, Y57, R99, Y101, and S106) and TMEM95 (P23, D26, S28, G29, R30, S36, S48, and R73). (c) Ribbon diagram of 6B08–TMEM95–FIMP, in which the 6B08 variable regions interact with the β-hairpin and α4′ helix of TMEM95. (d) Close-up view of the 6B08–TMEM95 interface. The interaction is mediated by hydrogen bonds and salt bridges between residues of the 6B08 heavy chain (V_H_: Y54, G55, T56, Y59, Y101, and D102) and TMEM95 (E102, R110, W129, and R131). The arrow indicates R110. (e) Biolayer interferometry traces showing binding of sensor-loaded TMEM95-Fc and TMEM95^R110A^-Fc to 200 nM 3A01 Fab or 6B08 Fab. Substitution of TMEM95 R110 with alanine does not affect 3A01 binding relative to wild-type TMEM95-Fc. (f) Model of 3A01-mediated disruption of the interaction of the TMEM95–FIMP complex with a neighboring protein. (g) Model of 6B08-mediated disruption of the orientation of the TMEM95–FIMP complex on the membrane.

3A01 recognized TMEM95 at the opposite surface from the FIMP-binding site, potentially blocking the interaction of TMEM95 with other sperm proteins (Fig. 3f and Extended Data Fig. 9a). 6B08 recognizes the membrane-proximal base of TMEM95, the surface below the FIMP-binding site. In the context of the sperm membrane, the bound 6B08 antibody would clash with the lipid bilayer, reorienting TMEM95–FIMP from an upright to a near-horizontal position and disrupting the functional architecture of the complex (Fig. 3g and Extended Data Fig. 9b). Together, these data provide a structural explanation for the activity of the fertilization-inhibitory, non-blocking antibodies 3A01 and 6B08 targeting the TMEM95–FIMP complex.

### The anti-TMEM95 antibody 3A02 recognizes the TMEM95–FIMP interface

We next examined whether the TMEM95–FIMP interaction is critical for fertilization. Deletion of *Tmem95* or *Fimp* in mice led to the loss of not only TMEM95 or FIMP, respectively, but also all known sperm fertilization proteins except IZUMO1 ^8,18,20^, making it difficult to interpret phenotypes specifically associated with the TMEM95–FIMP complex. To overcome this challenge and to biochemically pinpoint whether the TMEM95–FIMP interface is essential for sperm–egg interaction, we sought to elicit an antibody capable of disrupting the TMEM95–FIMP complex. Using a heterologous prime–boost strategy, we alternated immunization of a female mouse with murine and human TMEM95–FIMP ectodomain complexes (Fig. 4a). We generated hybridomas and screened for antibody cross-reactivity against human and murine TMEM95 (Fig. 4b and Extended Data Fig. 5a,b). As a result, we isolated 3A02 as a cross-reactive monoclonal antibody, a murine IgM with a kappa light chain. 3A02 belongs to the lineages carrying IGHV5-9-3*01 and IGKV3-4*01 gene segments, with low somatic mutations showing 97.6% and 99.3% identities to the heavy and light chain germline sequences, respectively (Extended Data Fig. 5c). We note minor sequence misalignments in the framework region 1 (FR1) of both chains likely reflecting gene amplification and sequencing errors rather than somatic mutations (Extended Data Fig. 5c).

**Fig. 4.**
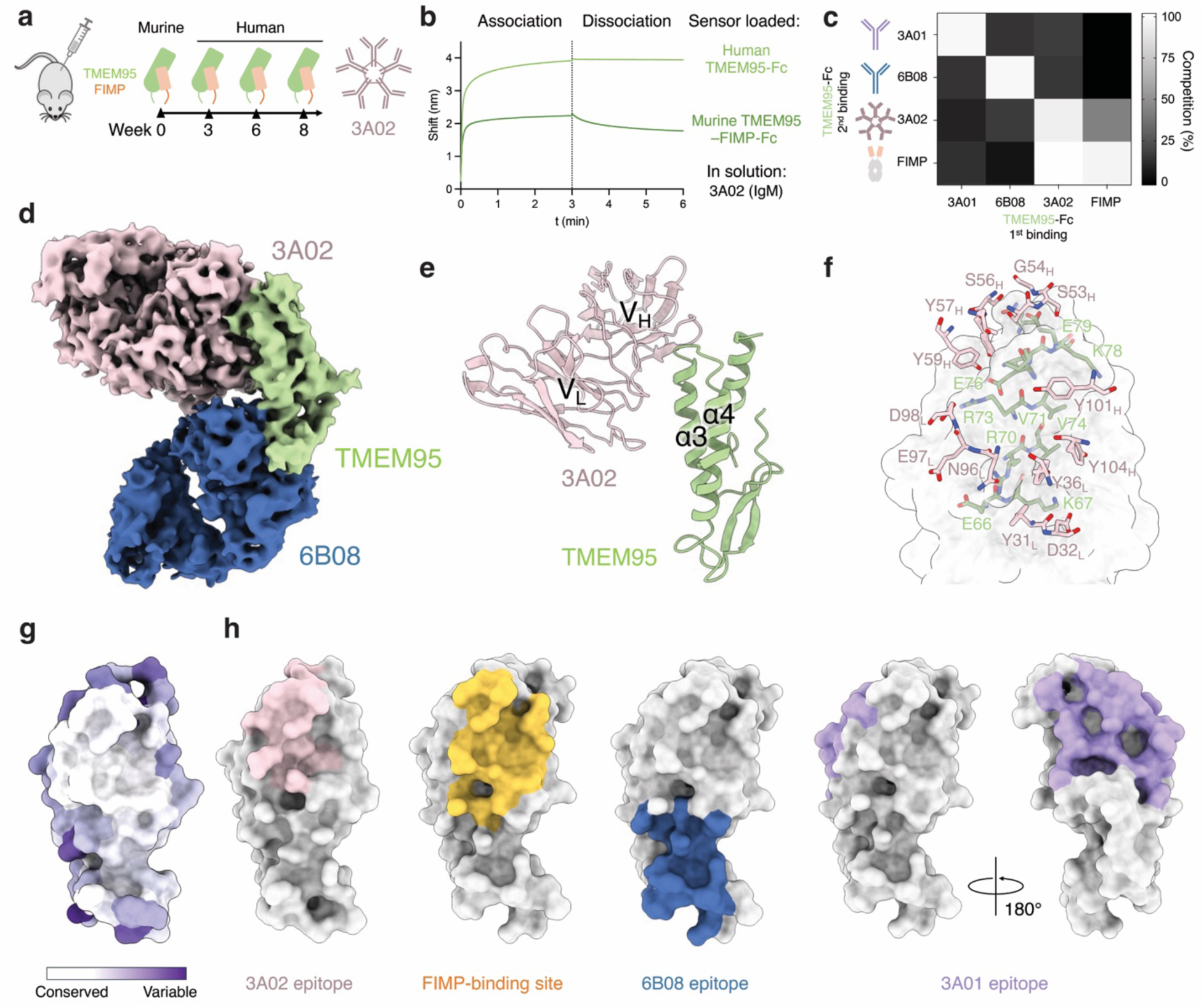
Structure of the human TMEM95–3A02–6B08 complex, showing that 3A02 recognizes the FIMP-binding site of TMEM95. (a) Schematic of mouse immunization: prime with murine TMEM95–FIMP proteins, boost with human TMEM95–FIMP proteins at weeks 3, 6, and 8, followed by hybridoma generation to isolate the 3A02 monoclonal antibody. (b) Biolayer interferometry traces showing binding of sensor-loaded human TMEM95-Fc (green) and murine TMEM95/FIMP-Fc (dark green) to 520 nM 3A02 (IgM). (c) Heat map summarizing competition between FIMP^F93A^-Fc, 3A01 IgG, 6B08 IgG, and 3A02 IgM for binding to sensor-immobilized TMEM95-Fc, showing competitive binding of FIMP^F93A^-Fc and 3A02 IgM to TMEM95. (d) Cryo-EM map of the human TMEM95–3A02–6B08 complex. TMEM95, green; 3A02, light pink; 6B08, dark blue. (e) Ribbon diagram of the cryo-EM structure of TMEM95–3A02, in which the 3A02 variable regions interact with the α3 and α4 helices of TMEM95. (f) Close-up view of the 3A02 variable region interacting with TMEM95. Residues S53, G54, and S56 of the 3A02 heavy chain (V_H_) form hydrogen bonds with I77, K78, E79, and Q37 of TMEM95, while D32, N96, and D98 of the 3A02 light chain (V_L_) form hydrogen bonds and salt bridges with K67, R70, and R73 of TMEM95. (g) Space-filling ConSurf model of TMEM95, with white indicating conserved regions and purple indicating variable regions among TMEM95 orthologs. (h) Space-filling model of the 3A02 epitope (light pink), FIMP-binding site (yellow), and 6B08 (dark blue) and 3A01 (purple) epitopes on the TMEM95 surface, showing overlapping footprints of the 3A02 epitope and FIMP-binding site on the conserved surface.

We found using BLI that 3A02 (IgM) can bind directly to the murine TMEM95–FIMP-Fc protein, or murine TMEM95-Fc “knobs” and murine FIMP-Fc “holes” protein, with a *K_D_,* _apparent_ of 20.74 ± 7.78 nM, suggesting that 3A02 (IgM) can effectively compete off murine FIMP to bind murine TMEM95 in a pre-formed heterodimeric murine TMEM95–FIMP complex (Fig. 4b). To further determine whether 3A02 blocks human TMEM95–FIMP complex formation, we performed a BLI-based competition analysis using human TMEM95-Fc to bind to 3A01 (IgG), 6B08 (IgG), 3A02 (IgM), and FIMP^F93A^-Fc. We found that while 3A02 does not compete with 3A01 or 6B08, pre-binding of 3A02 to TMEM95-Fc effectively blocked subsequent FIMP^F93A^-Fc binding. Moreover, when FIMP^F93A^-Fc was pre-bound to TMEM95-Fc, 3A02 displaced approximately 50% of bound FIMP^F93A^-Fc (Fig. 4c), indicating that 3A02 can substantially disrupt the pre-formed human TMEM95–FIMP complex. Together, these results define three non-overlapping epitopes on TMEM95 recognized by 3A01, 6B08, and 3A02, respectively, with the 3A02 epitope overlapping with the FIMP-binding site.

To characterize how 3A02 binds TMEM95, we assembled a ternary complex of human TMEM95 ectodomain, 6B08 Fab, and 3A02 Fab (Extended Data Fig. 5d,e) and determined a cryo-EM structure of the complex at 3.9 Å resolution (Fig. 4d, Extended Data Fig. 6a,b,c,d,e,f, Table 1, and Extended Data Table 2.) In this structure, TMEM95, 6B08, and 3A02 are present in a 1:1:2 stoichiometry, with two 3A02 Fabs forming an anti-parallel homodimer (Extended Data Fig. 7a,b,c). The dimeric 3A02 Fabs are consistent with their early retention volume in size-exclusion chromatography (Extended Data Fig. 5e) compared to a monomeric Fab (Extended Data Fig. 2f). Only one 3A02 Fab protomer contacts TMEM95, where CDRs from the heavy and light chains interact with the α3 and α4 helices of TMEM95 (Fig. 4e,f). The epitope of 3A02 overlaps with the FIMP-binding site, the conserved surface on TMEM95 (Fig. 4g,h and Extended Data Fig. 7d,e,f), providing a structural basis for 3A02’s cross reactivity. These biochemical properties of 3A02 enable the use of this antibody to investigate the functional role of the TMEM95–FIMP interaction in mouse fertilization.

### The anti-TMEM95 antibody 3A02 impairs murine sperm–egg fusion

We next examined whether the TMEM95–FIMP complex is essential for fertilization in mice. The ∼25 kDa single-chain fragment variable (scFv) of 3A02 binds TMEM95 with less avidity and may have less steric effects in membrane fusion than its larger, ∼900 kDa IgM counterpart (Fig. 5a). We produced the IgM and the scFv of the anti-TMEM95 3A02 antibody (Extended Data Fig. 7g) and tested them in a sperm–egg fusion assay using wild-type mice. We incubated zona-free eggs with sperm at 2 × 10^4^ sperm per mL pretreated with the antibodies (Fig. 5b and Extended Data Fig. 8a). We used an untreated group as a negative control and anti-IZUMO1 antibody OBF13 (IgM) and OBF13^HAC^ (scFv) as positive controls. The OBF13 antibodies were previously shown to block acrosome-reacted sperm from binding to eggs, thereby preventing membrane fusion ^21^. At 6 h after insemination, we recorded the number of pronuclei per egg to evaluate fusion. In the untreated group, there were approximately 1.5 ± 0.23 (mean ± standard error of the mean) fused sperm per egg, and in the OBF13 (IgM)- and OBF13^HAC^ (scFv)-treated groups, no sperm–egg fusion was detected. The average numbers of fused sperm per egg significantly decreased to 0.5 ± 0.19 (*P* = 0.0032) and 0.5 ± 0.23 (*P* = 0.0056) in the anti-TMEM95 3A02 IgM and scFv groups, respectively (Fig. 5c). When we increased the sperm concentration to 2 × 10^5^ cells per mL, we observed a similar inhibitory effect in the sperm–egg fusion assay for the anti-TMEM95 3A02 IgM and scFv groups (Extended Data Fig. 8b,c,d). Together, the anti-TMEM95 3A02 antibody that targets the FIMP-binding site of TMEM95 impairs sperm–egg fusion, suggesting that the TMEM95–FIMP interface plays a role in sperm–egg interaction (Fig. 5d and Extended Data Fig. 9d).

**Fig. 5.**
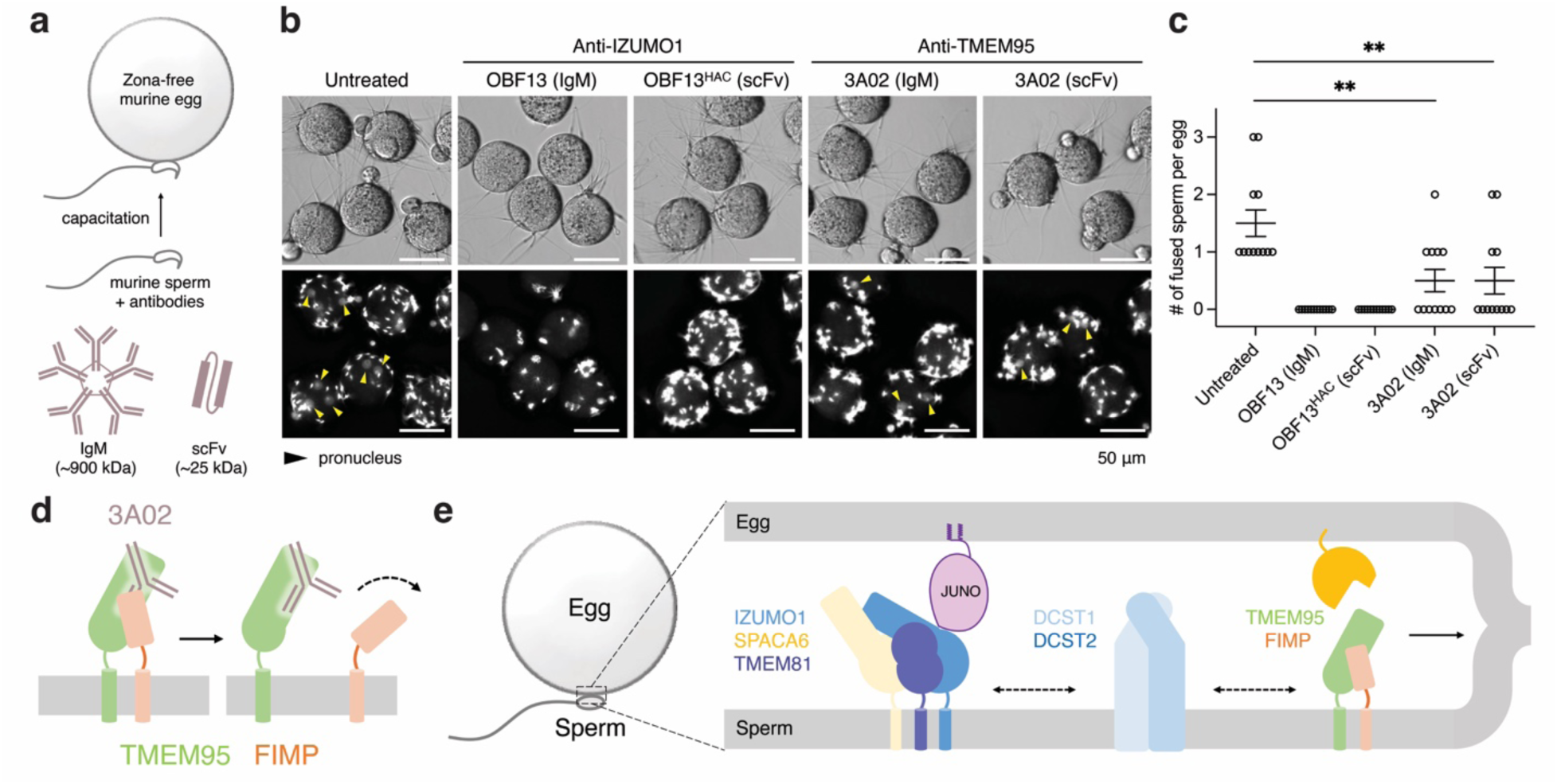
The anti-TMEM95 antibody 3A02 inhibits murine sperm–egg fusion *in vitro*. (a) Cartoon schematic of the in vitro fertilization experiment using wild-type mouse sperm and eggs. The zona pellucida was removed to yield zona-free eggs. Sperm were treated with antibodies, capacitated, and incubated with zona-free eggs. (b) Representative micrographs of sperm and zona-free eggs 6 hours after insemination. DNA was labeled with Hoechst 33342. Sperm were untreated or pretreated with 50 μg/mL OBF13 (IgM), 50 μg/mL OBF13^HAC^ (scFv), 440 μg/mL 3A02 (IgM), or 50 μg/mL 3A02 (scFv). Scale bars, 50 μm. (c) Number of fused sperm per zona-free egg (mean ± standard error of the mean, SEM): untreated, 1.5 ± 0.23 (*N* = 12); OBF13 (IgM)-treated, 0 ± 0 (*N* = 12); OBF13^HAC^ (scFv)-treated, 0 ± 0 (*N* = 12); 3A02 (IgM)-treated, 0.5 ± 0.19 (*N* = 12); 3A02 (scFv)-treated, 0.5 ± 0.23 (*N* = 12). (d) Model of 3A02-mediated disruption of the TMEM95–FIMP complex. Despite the experimental use of IgM and scFv, a Y-shaped IgG is used to depict 3A02 as an antibody. (e) Model of mammalian sperm–egg interaction. Schematic showing three sperm membrane protein complexes in the fertilization synapse: the IZUMO1–TMEM81–SPACA6 complex, the DCST1–DCST2 complex, and the TMEM95–FIMP complex. Dashed arrows indicate potential interactions among complexes; the solid arrow indicates that a yet-to-be-identified mechanism drives sperm–egg membrane fusion.

## Discussion

Two fertilization-essential sperm acrosomal membrane proteins, TMEM95 and FIMP, form a high-affinity complex through their ectodomains. The previously identified, evolutionarily conserved surface of TMEM95 interacts with an aromatic motif of FIMP. A near-germline antisperm antibody isolated in this study, 3A02, targets this conserved surface and inhibits membrane fusion during fertilization. Together with the IZUMO1–SPACA6–TMEM81 and DCST1–DCST2 complexes, the TMEM95–FIMP complex plays an essential role in sperm–egg interaction at the mammalian fertilization synapse (Fig. 5e).

The human TMEM95–FIMP ectodomain structure reported here reveals that a conserved FFDY motif of FIMP interacts with TMEM95 at the putative egg-receptor-binding site (Fig. 2f,g,h and Extended Data Fig. 4d). The anti-TMEM95 antibody 3A02, isolated in this study, also binds to this site. 3A02 is fertilization-inhibitory, potentially by disrupting both the FIMP-mediated and egg-receptor interactions of TMEM95. One possibility is that the TMEM95–FIMP complex functions as a heterodimer during membrane fusion. Alternatively, or in addition, FIMP may be displaced from TMEM95 upon sperm–egg interaction, making the surface of TMEM95 available for interacting with the egg. Evidence supporting this notion includes the observation that FIMP becomes undetectable in ∼40% of acrosome-reacted murine sperm ^2^, and that TMEM95 predates FIMP evolutionarily, with TMEM95 orthologs supporting fertilization in some vertebrates that lack FIMP (Extended Data Fig. S10a,b,c). Future structure-based engineering of 3A02 through directed evolution or protein language model-guided design could yield high-affinity variants ^21,22^, which could serve as tools to capture new conformational intermediates of TMEM95 during sperm–egg interaction.

The FIMP-mediated interaction of TMEM95 identified here also provides a structural basis for the discovery and design of contraceptives. First, a pocket that forms on the conserved surface of TMEM95 upon FIMP binding may represent an attractive small-molecule contraceptive target across diverse mammalian species (Extended Data Fig. 4d,f). The structures reported here will facilitate virtual drug screening to identify potential lead compounds. Specifically, we envision a small molecule that contacts all or many of the residues lining the pocket, particularly K67, R70, V71, V74, Y87, and W90, in a conformation similar to that formed in the complex with FIMP (Extended Data Fig. 4f). In addition, the benzyl and phenol rings and the neighboring side chains of the FFDY motif of FIMP may serve as useful starting points for fragment-based screening scaffolds (Extended Data Fig. 4f). Second, the discovery of 3A02 suggests avenues for anti-TMEM95 immunocontraceptives. An inferred germline precursor of 3A02 binds TMEM95 (Extended Data Fig. 7g) with an affinity above the threshold required for B cell activation and affinity maturation ^21,23,24^. The murine adaptive immune system can elicit antisperm antibodies against syngeneic sperm targeting this cryptic but immunogenic site on TMEM95.

Anti-TMEM95 antibodies 3A01 and 6B08 do not block the FIMP-mediated interaction of TMEM95 but inhibit membrane fusion of human sperm with hamster eggs ^7^. How could these non-FIMP-blocking antibodies inhibit membrane fusion? Evidence suggests that TMEM95–FIMP is part of a hetero-oligomeric sperm membrane protein complex ^18^. Two evolutionarily conserved, solvent-accessible patches outside the TMEM95–FIMP interface may serve as interfaces for additional sperm fertilization proteins (Extended Data Fig. 9a). One lies immediately adjacent to the 3A01 epitope, implying that 3A01 may inhibit fertilization by sterically interfering with interactions between TMEM95–FIMP and additional sperm proteins (Extended Data Fig. 9a). Overlay of our structure of the 3A01- and 6B08-bound TMEM95–FIMP complex with an AlphaFold3 model of the larger sperm complex showed that 3A01 and the DCST1–DCST2 complex would clash if 3A01 were to bind TMEM95 in its hetero-oligomeric state, separating TMEM95–FIMP from the rest of the complex (Extended Data Fig. 9b). Binding of 6B08 to the membrane-proximal base of TMEM95 would clash with the sperm membrane, potentially displacing TMEM95 from its native membrane-bound conformation. Such perturbation may reorient TMEM95 and disrupt its hetero-oligomeric assembly and its binding to the egg (Extended Data Fig. 9c). Both antibodies recognize TMEM95 at nanomolar affinities predominantly through their heavy chains, making them suitable candidates for single-chain nanobody engineering to enable conformation-specific recognition and inhibition of TMEM95.

Taken together, we propose that mammalian sperm–egg interaction involves three identified sperm membrane protein complexes: IZUMO1–SPACA6–TMEM81, DCST1–DCST2, and TMEM95–FIMP (Extended Data Fig. 10a,b,c). The recently identified acrosomal membrane proteins TMDD1 and FAM187A are essential for fertilization and might bridge these complexes on the sperm membrane to coordinate their function ^18,19^. When these sperm complexes assemble during spermatogenesis and sperm–egg interaction remains an open question. We anticipate additional analogous, yet-to-be-identified, interactions between sperm proteins and their egg receptors. More broadly, our work takes steps toward fully understanding the molecular interactions of the mammalian fertilization complex and has direct implications for the design and development of contraceptives.

## Materials and methods

### AlphaFold3 screen for testis-enriched proteins that interact with TMEM95

The ectodomain of human TMEM95 (residues 17–138) was used as bait. The N-linked glycan N-acetylglucosamine (NAG) was modeled as a ligand covalently bonded to the side-chain nitrogen of N117. A library of protein targets was generated from the Human Protein Atlas (HPA, v25.proteinatlas.org), filtered for proteins expressed in testis (mRNA level) and annotated by the HPA “protein class” field as predicted membrane or secreted soluble proteins. Custom R (version 4.4.1) and command-line scripts were used to generate AlphaFold3 input files.

AlphaFold3 (version 3.0.1) predictions were run with default settings. Pairwise ipTM and minimum PAE scores, averaged across the five models per bait–target complex, were used to rank and evaluate predictions. Structures were visualized with ChimeraX and PAE Viewer.

### Protein expression and purification

The TMEM95-Fc “knobs” expression construct encodes the human TMEM95 ectodomain (residues 17–138) fused to Fc with two substitutions, S237C and T249W ^25^. The FIMP F93A-Fc holes expression construct encodes the human FIMP ectodomain (residues 58–97) with the F93A substitution fused to Fc with four substitutions, T232C, T249S, L251A, and Y290V ^25^. For protein expression, plasmids encoding TMEM95-Fc “knobs”, FIMP F93A-Fc “holes”, anti-TMEM95 6B08 Fab heavy chain with a Ctag, and anti-TMEM95 6B08 light chain were co-transfected into HEK293F cells. The TMEM95–FIMP-Fc–6B08 Fab complex was purified from conditioned media using a CaptureSelect CtagXL affinity matrix (ThermoFisher), followed by size-exclusion chromatography on a Superdex 200 Increase column (GE Life Sciences) in HBS (150 mM NaCl, 20 mM HEPES pH 7.4).

To obtain the TMEM95–6B08 Fab complex, plasmids encoding the TMEM95 ectodomain, anti-TMEM95 6B08 heavy chain, and 6B08 light chain were co-transfected into HEK293F cells. The TMEM95–6B08 Fab complex, the human TMEM95–FIMP-Ctag complex, and the mouse TMEM95 (residues 17–141)–FIMP (residues 39–84)-Ctag complex were purified using approaches as described above.

To obtain the 3A01 and 3A02 Fab fragments, plasmids encoding the heavy and light chains were co-transfected into HEK293F cells. The Fab fragments were purified from conditioned media on a HiTrap Protein G HP column (Cytiva), followed by size-exclusion chromatography on a Superdex 200 Increase column in HBS.

To obtain OBF13 (IgM) and 3A02 (IgM), hybridoma culture supernatants were harvested and the IgM antibodies were purified using a LigaTrap™ (Dianova) column, followed by size-exclusion chromatography on a Superose 6 Increase column in HBS.

To obtain the 3A02 and 3A02 germline-reverted scFv fragments, plasmids encoding the heavy- and light-chain variable regions joined by a 19-residue GS linker, with a C-terminal 6×His tag, were transfected into HEK293F cells. The scFv fragments were purified from conditioned media on Ni-NTA resin (Invitrogen) and then on a Superdex 200 Increase column in HBS. The Fc fusion proteins (TMEM95-Fc, FIMP-Fc, TMEM95–FIMP-Fc, murine FIMP-Fc, and murine TMEM95–FIMP-Fc) were purified by the same methods.

### Flow Cytometry of HEK293F cell-surface binding

HEK293F cells were transiently transfected with a plasmid encoding full-length mouse TMEM95. Two days post-transfection, cells were collected by centrifugation, washed, and resuspended in ∼100 μL of selection buffer (150 mM NaCl, 20 mM HEPES pH 7.4, 0.5% BSA). An aliquot of 2.5 μL of the cell suspension was incubated with 100 μL of serially diluted murine FIMP-Fc for 1 hour at room temperature. Following incubation, cells were washed with 1 mL of selection buffer and resuspended in 100 μL of the same buffer containing a 1:500 dilution of either anti-human Fc-Alexa Fluor 647 or anti-mouse Fc-Alexa Fluor 647 secondary antibody and incubated for 20 minutes. Cells were then washed again with 1 mL of buffer and resuspended in 100 μL of selection buffer. Flow cytometry was performed using a BD Accuri™ C6 flow cytometer, and 1,000–10,000 events were collected per sample. Events were recorded and measured in the FL4 channel and analyzed to generate binding curves using GraphPad Prism 11.

### Cryo-EM data collection, processing, and model building

To prepare the human TMEM95–FIMP-Fc–3A01–6B08 sample for cryo-EM structure determination, the human TMEM95–FIMP-Fc–6B08 complex was mixed with 3A01 Fab at a 1:1.5 molar ratio and incubated overnight at a final concentration of 1 mg/mL. 4 μL of the complex was applied to a glow-discharged holey-carbon gold grid (Quantifoil R1.2/1.3) in an FEI Vitrobot Mark IV (4 °C, 100% humidity). Grids were blotted for 3 s after a 0.5-s wait and plunged into liquid ethane. Grids were screened on a Titan Krios operating at 300 kV equipped with a K3 direct electron detector (Gatan), and one grid was selected for data collection. Movie stacks were collected automatically at a calibrated magnification of 105,000× using EPU, over a defocus range of −1.0 to −2.5 μm. Each micrograph was recorded with a total electron dose of ∼50 e⁻/Å² at an exposure rate of 15 e⁻/Å²/s, fractionated into 50 frames. An energy filter with a 20-eV slit was used during data collection.

To prepare the human TMEM95–3A02–6B08 sample for cryo-EM structure determination, the human TMEM95-Ctag–6B08 complex was mixed with 3A02 Fab at a 1:1.5 molar ratio and incubated overnight at a final concentration of 1 mg/mL. Sample preparation and data collection were as described above.

All image processing was performed in cryoSPARC. Drift correction and electron-dose weighting were applied to movie stacks using the Patch Motion Correction (multi) job. Contrast transfer function (CTF) parameters were estimated for each summed image using the Patch CTF Estimation (multi) job. Micrographs were screened manually with the Manually Curate Exposures job, and qualified micrographs were selected for further processing. Particles were picked using Blob Picker, and extracted particles were subjected to reference-free 2D classification to remove noise and junk particles. Selected particles were used for ab initio reconstruction and heterogeneous refinement. Particles from qualified classes were then pooled and subjected to homogeneous refinement. Model building and refinement were performed in Coot and Phenix. Structural figures were prepared in UCSF ChimeraX.

### Mouse hybridoma antibody generation and sequencing

Five female BALB/c mice (Jackson Laboratory) aged ∼8 weeks were immunized with 10 μg purified protein of murine TMEM95–FIMP-Ctag in 100 μL of 150 mM NaCl, 20 mM HEPES pH 7.4, adjuvanted with 10 μg Quil-A (InvivoGen) and 10 μg monophosphoryl lipid A (InvivoGen). Mice were boosted with human TMEM95–FIMP-Ctag at days 21, 33, 41, and 58. At day 61, a spleen of one mouse was disaggregated into a single-cell suspension for hybridoma generation following the manufacturer’s procedures (STEMCELL technologies). This work was approved by Yale University’s Institutional Animal Care and Use Committee (IACUC, 2024-20562). The conditioned media of hybridomas were screened for binding to TMEM95 or FIMP by enzyme-linked immunosorbent assay (ELISA) and biolayer interferometry (BLI). Binding-positive clones were expanded for purification and antibody sequencing ^26^.

Total RNA was isolated using TRIzol reagent (Invitrogen™) from 5 mL hybridoma cells following the manufacturer’s instructions. The cDNA was generated by reverse transcription (ProtoScript® First Strand cDNA Synthesis Kit). The variable regions of immunoglobulin heavy (VH) and light (VL) chains were amplified by PCR using a set of degenerate primers specific for murine IgG variable region frameworks. The PCR amplification primers were used as previously described ^26^. PCR products were purified and cloned into a pVRC expression vector for Sanger sequencing.

### Biolayer interferometry (BLI)

An Octet RED96 system (Sartorius) was employed for protein–protein interaction assays. Biotinylated TMEM95-Fc proteins were loaded onto Streptavidin biosensors (Sartorius), the protein to be tested was diluted to different concentrations in Octet buffer (150 mM NaCl, 20 mM HEPES pH 7.4, 0.1% bovine serum albumin, and 0.05% Tween 20). Loaded biosensors were equilibrated in Octet buffer (baseline), dipped into analyte proteins diluted to a range of concentrations in Octet buffer (association), and returned to Octet buffer (dissociation). Baseline correction was performed in Octet Analysis Studio 13.0.3.52, and binding curves were plotted in GraphPad Prism 11. All experiments were performed with three biological replicates. Data are presented as mean ± standard deviation.

### Epitope mapping by biolayer interferometry competition assay

BLI competition experiments were performed on an Octet system (Sartorius). Biotinylated TMEM95-Fc was immobilized onto Streptavidin biosensors (Sartorius) to a response of 0.5 nm and equilibrated in Octet buffer. Loaded biosensors were saturated with a first binding partner, FIMP-Fc, 3A01 IgG, 6B08 IgG, or 3A02 IgM, for 480 s, returned briefly to Octet buffer for a baseline step, dipped into a second binding partner (FIMP-Fc, 3A01 IgG, 6B08 IgG, or 3A02 IgM) for association, and returned to Octet buffer for dissociation.

Association and dissociation signals were recorded in real time at 25 °C. Data were reference-subtracted and analyzed in Octet Analysis Studio 13.0.3.52. Competition was quantified using the binding response (nm) at 120 s of the second-partner association step. The response in the presence of the first partner was divided by the response of the second partner alone under the same conditions, and this ratio was used as a measure of competition between the two partners. The competition heatmap was plotted in GraphPad Prism 11.

### Conservation analysis

Homologous sequences were collected by NCBI BLAST and aligned with a multiple sequence alignment. Per-residue evolutionary conservation scores were calculated with the ConSurf server ^27^. Sequence conservation was also analyzed using the Sequence Conservation function in UCSF ChimeraX. Conservation scores were mapped onto the protein structure and visualized in UCSF ChimeraX, with conserved residues shown in white/purple and variable residues in darker colors, as indicated in the figure legends.

### Sperm–egg fusion assay using *in vitro* fertilization

B6D2F1 mice, used as wild-type mice in this study, were either purchased from Japan SLC, Inc. or bred in-house by pairing C57BL/6 females with DBA/2 males. Mice were maintained under a 14 h light/10 h dark cycle with ad libitum access to food and water. All animal experiments were approved by the Institutional Animal Care and Use Committee of The University of Osaka Medical School (#24-020-007) and were conducted in compliance with all applicable guidelines and regulations.

Cauda epididymal sperm collected from 10- to 12-week-old wild-type males were preincubated in Toyoda–Yokoyama–Hosi (TYH) medium drops with or without antibodies at a density of 2 × 10^4^ or 2 × 10^5^ sperm/mL for 2 h to induce capacitation. Eggs obtained from 6-week-old superovulated wild-type females were treated with 1 mg/mL of collagenase from *Clostridium histolyticum* (Sigma Aldrich) for 10 min to remove both the cumulus cells and the zona pellucida, followed by staining with 1:1,000 Hoechst 33342 for 15 min. The treated eggs were then washed in fresh TYH medium drops and transferred to the sperm drops. After 6 h of insemination, the number of pronuclei in each egg was counted and z-stack images were captured under a Keyence microscope. All incubation steps were performed at 37 °C under 5% CO_2_.

## Data availability

The cryo-EM maps of the human TMEM95–FIMP–3A01–6B08 complex and the human TMEM95–6B08–3A02 complex, along with their corresponding atomic coordinates, have been deposited in the Electron Microscopy Data Bank (EMDB) and the Protein Data Bank (PDB) under the accession codes EMD-77083 and 13IN, and EMD-77090 and 13JC, respectively. All data analyzed in this study are included in this paper. Other relevant data are available from the corresponding author upon reasonable request.

## Code availability

The AlphaFold3 screen scripts are available at https://github.com/rcastelino01/AlphaScreen.

## Acknowledgements

We thank Drs. Andrea Pauli, Masahito Ikawa, and Amber Krauchunas for discussion and for coordinating our manuscripts. We thank the Yale Center for Research Computing (YCRC) for computational resources, and Dr. Patrick Mitchell at the Stanford-SLAC Cryo-EM Center (S2C2) for supporting our cryo-EM data collection; S2C2 is supported by NIH NIGMS grant 1R24GM154186. We thank Kaori Nishino, Yufei Wang, Dr. Takemasa Kawashima, Dr. Chengtao Xu, and Dr. Tianying Qiu for technical support. This work was supported by the NSF Graduate Research Fellowship Program (DGE-2139841 to T.G.); the JSPS Grant-in-Aid for Scientific Research (B) (JP24K02033 to Y.L.); the Takeda Science Foundation Life Science Research Grant (2024032851 to Y.L.); the Asahi Glass Foundation Research Encouragement Grant (Y.L.); the PRIMe Joint Research Grant (1301800004 to Y.L.); the Mochida Memorial Foundation for Medical and Pharmaceutical Research (Y.L.); NIH NICHD grant (R00HD104924 to S.T.); and the David Sokal Innovation Award of the Male Contraceptive Initiative (2024-303 to S.T.).

## Author contributions

P.L. and S.T. designed and performed most of the experiments. R.E.C. performed the AlphaFold3 screen. Y.L. performed the *in vitro* fertilization experiments. S.T. conceived and supervised the project. All authors contributed to drafting the manuscript.

---

**Extended Data Fig. 1.**
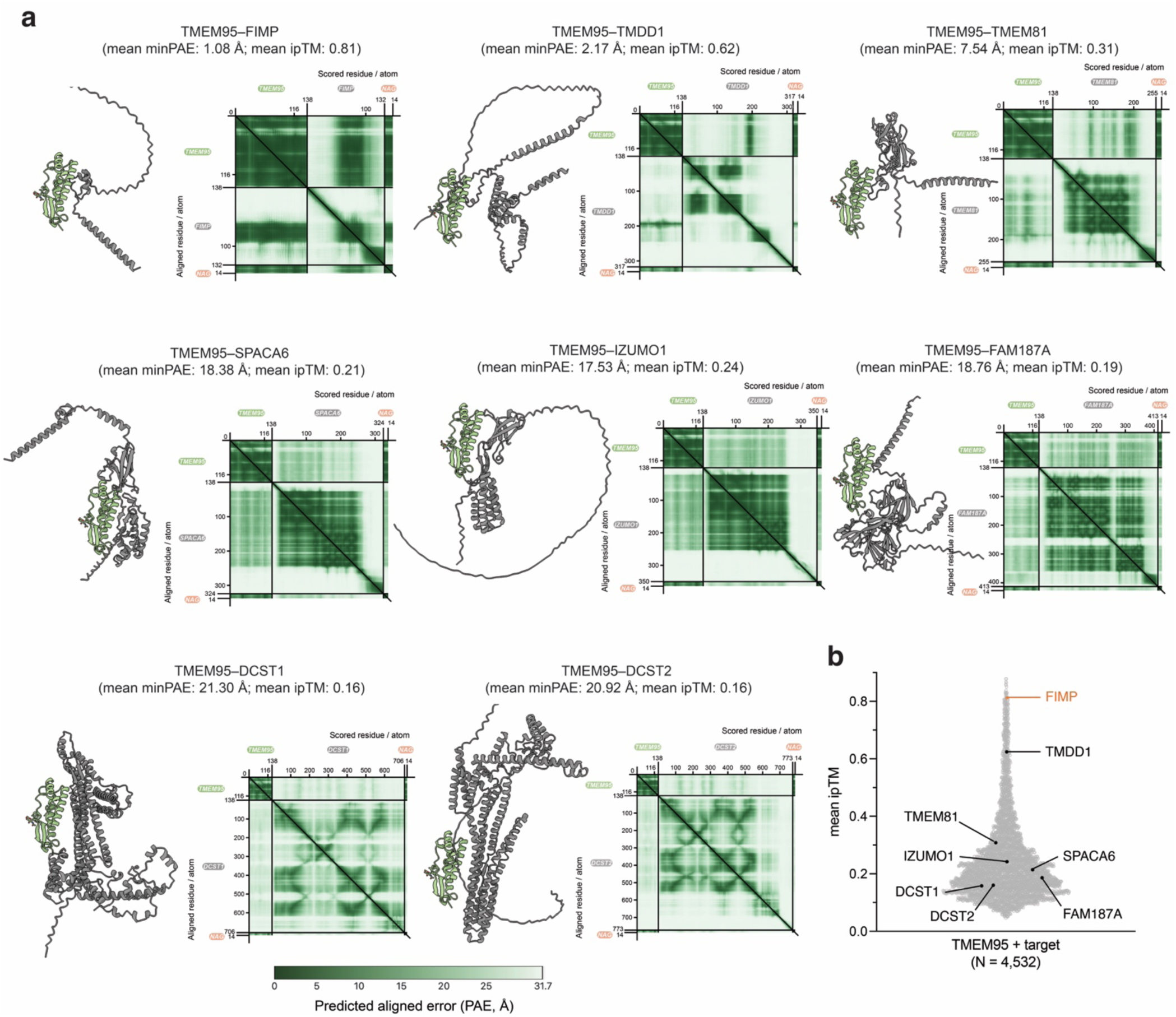
AlphaFold3-predicted structures of TMEM95 in complex with sperm fertilization proteins. (a) AlphaFold3 predictions of TMEM95’s ectodomain (residues 17–138), shown in light green, in complex with known fertilization factors FIMP, TMDD1, TMEM81, SPACA6, IZUMO1, FAM187A, DCST1, and DCST2, shown in grey. The conserved N-glycan of TMEM95, N-acetylglucosamine (NAG), is shown as sticks. Predicted aligned error (PAE) plots for each predicted complex are shown, with dark green indicating lower predicted aligned error and higher confidence in relative residue positioning. (b) Violin plot of ipTM scores averaged across the five predicted models. FIMP is labeled in yellow; other fertilization factors are labeled in black.

**Extended Data Fig. 2.**
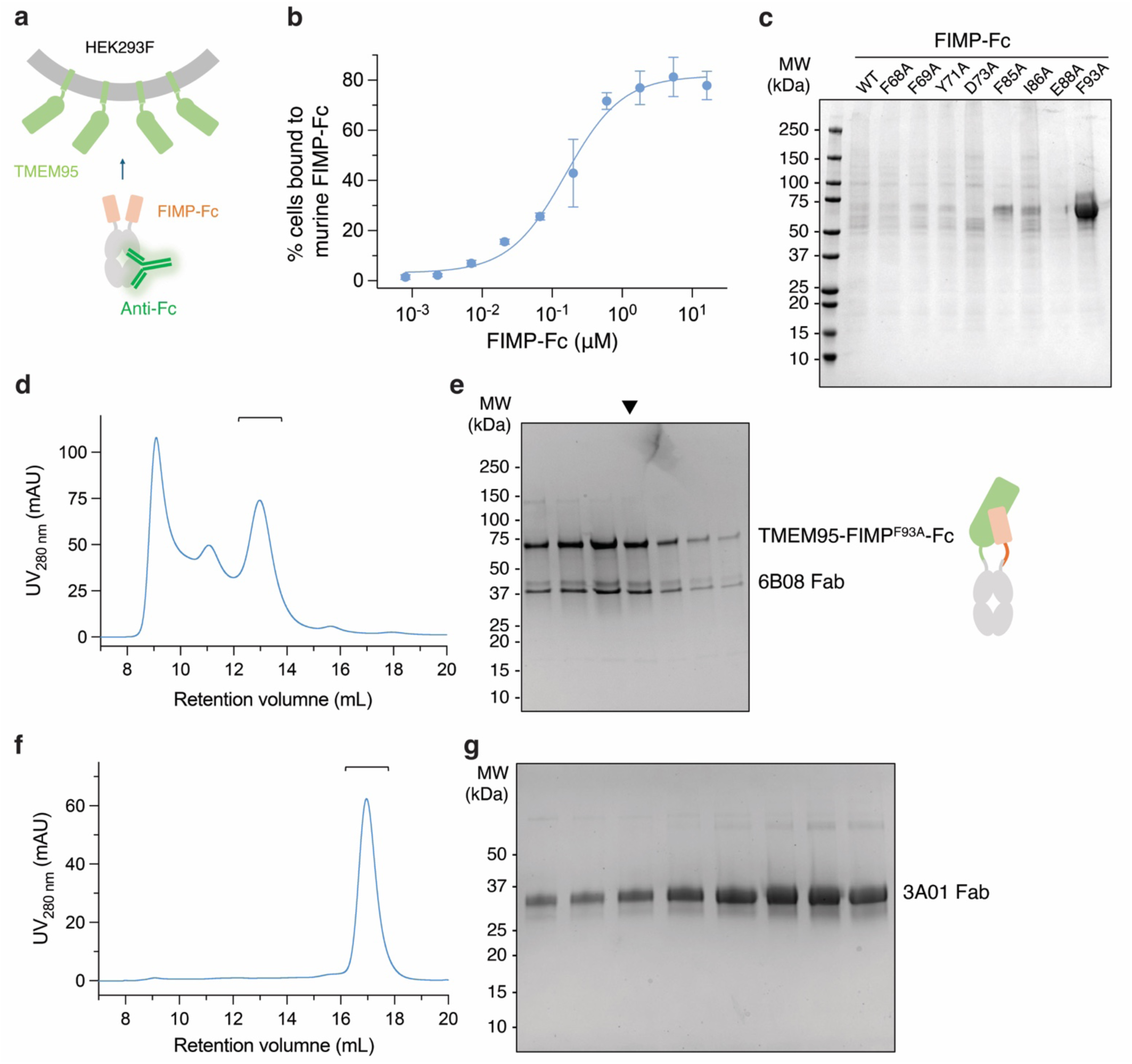
Biochemical purification and characterization of the TMEM95–FIMP–3A01–6B08 complex. (a) Cartoon schematic of (b) the flow cytometry experiment using HEK293F cells expressing full-length murine TMEM95 labeled with a serial dilution of murine FIMP-Fc and an AlexaFluor647-conjugated anti-Fc antibody. (c) Coomassie-stained SDS–PAGE of FIMP-Fc variant proteins. Single amino acid substitutions (F85A, I86A, or F93A) each dramatically increased protein expression levels compared to wild-type FIMP-Fc. (d) Size-exclusion chromatogram of TMEM95–FIMP–6B08 Fab, showing three distinct peaks. (e) Peak fractions analyzed by SDS–PAGE, showing co-elution of TMEM95–FIMP-Fc with 6B08 Fab. The fraction marked with a triangle was used for cryo-EM sample preparation. (f) Size-exclusion chromatogram and (g) Coomassie-stained SDS–PAGE of 3A01 Fab used for cryo-EM sample preparation.

**Extended Data Fig. 3.**
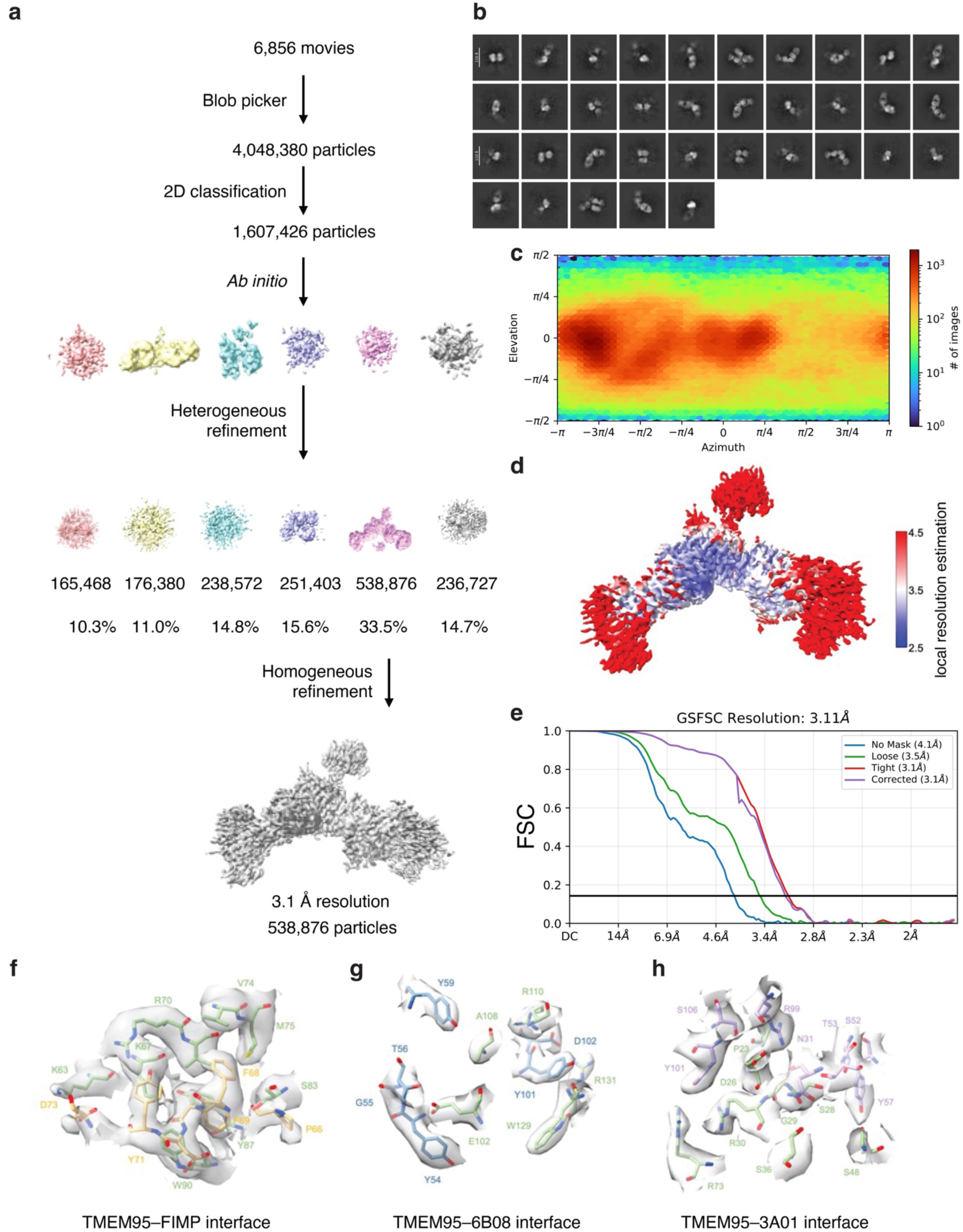
Cryo-EM data processing workflow of the human TMEM95–FIMP–3A01–6B08 complex. (a) Flow chart of cryo-EM data processing. (b) Representative 2D class averages. Scale bar, 110 Å. (c) Eulerian angle distribution of the particles used in the final 3D reconstruction. (d) Local resolution estimation of the density map. (e) Gold-standard Fourier shell correlation (FSC) curve with the estimated resolution for the overall density map. (f) Local cryo-EM density of the TMEM95–FIMP interface. TMEM95 residues are shown in green; FIMP residues in yellow. Local cryo-EM density of (g) the TMEM95–6B08 interface and (h) the TMEM95–3A01 interface.

**Extended Data Fig. 4.**
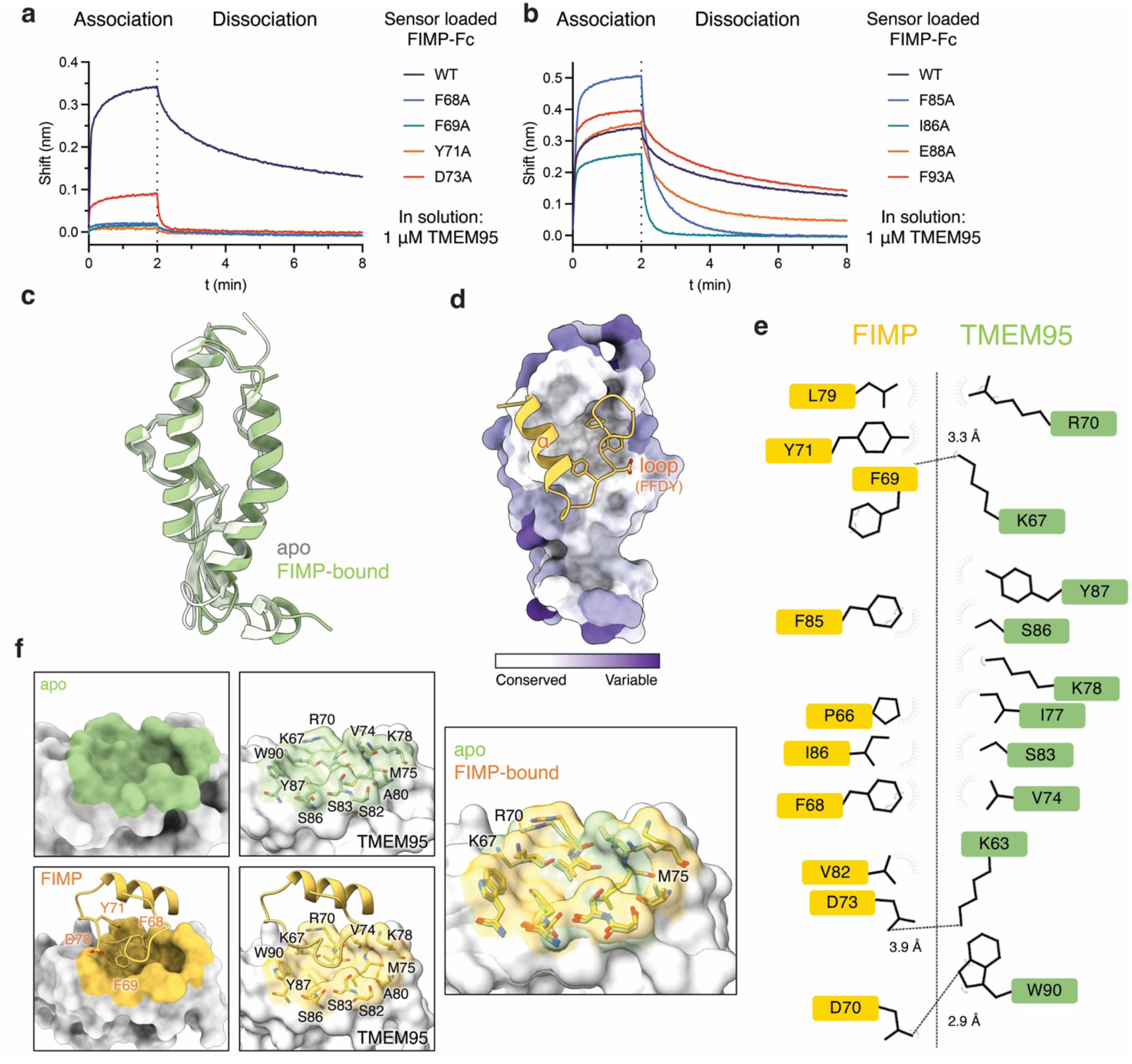
The TMEM95–FIMP interaction and the conformations of the FIMP-binding pocket on TMEM95. Bio-layer interferometry traces showing binding of sensor-loaded (a) wild-type (WT), F68A, F69A, Y71A, or D73A FIMP-Fc variants or (b) wild-type (WT), F85A, I86A, E88A, or F93A FIMP-Fc variants to 1 μM TMEM95. Association, 2 min; dissociation, 6 min. (c) Structural overlay of TMEM95 in the apo (light green) and FIMP-bound (green) states. (d) FIMP (yellow ribbon) binds to a conserved surface on TMEM95 (surface representation). The FFDY motif of FIMP is shown in stick representation. Conservation scores were mapped onto TMEM95, with residues colored from white (conserved) to purple (variable). (e) 2D schematic of the interactions between TMEM95 (green) and FIMP (yellow). Hydrogen bonding interactions are shown as dashed lines and van der Waals forces are depicted as grey semi-circles. (f) Space-filling models of the FIMP-binding site of TMEM95 in (upper) the apo state or (bottom) the FIMP-bound state. The FFDY motif of FIMP binds into a defined pocket on TMEM95. Structural overlay of apo and FIMP-bound TMEM95 (right), illustrating conformational differences in pocket-forming residues, especially K67, R70, and M75.

**Extended Data Fig. 5.**
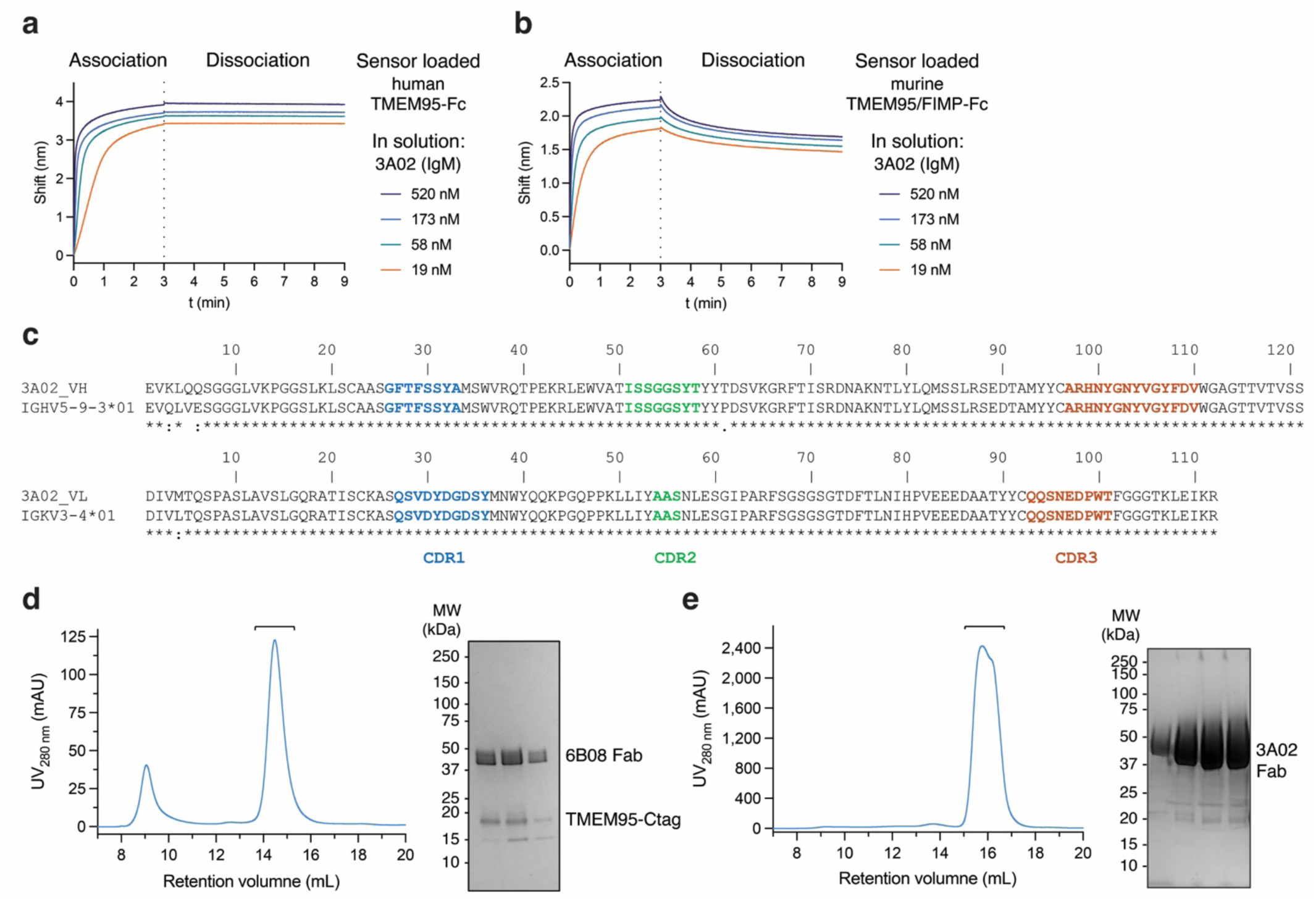
Characterization and purification of the anti-TMEM95 3A02 Fab. Biolayer interferometry traces showing (a) sensor-loaded TMEM95-Fc or (b) sensor-loaded murine TMEM95–FIMP-Fc binding to 520 nM, 173 nM, 58 nM, or 19 nM 3A02 (IgM). Association, 3 min; dissociation, 6 min. (c) Sequence alignment of 3A02 heavy chain variable region with its inferred germline sequence, and of 3A02 light chain variable region with its inferred germline sequence; the three complementarity-determining regions (CDRs) are highlighted. (d) Size-exclusion chromatogram of TMEM95 in complex with 6B08 Fab, showing two peaks, and Coomassie-stained SDS–PAGE of fractions from the second peak, which was used for cryo-EM sample preparation. (e) Size-exclusion chromatogram of 3A02 Fab and Coomassie-stained SDS–PAGE of fractions from the peak, which was used for cryo-EM sample preparation.

**Extended Data Fig. 6.**
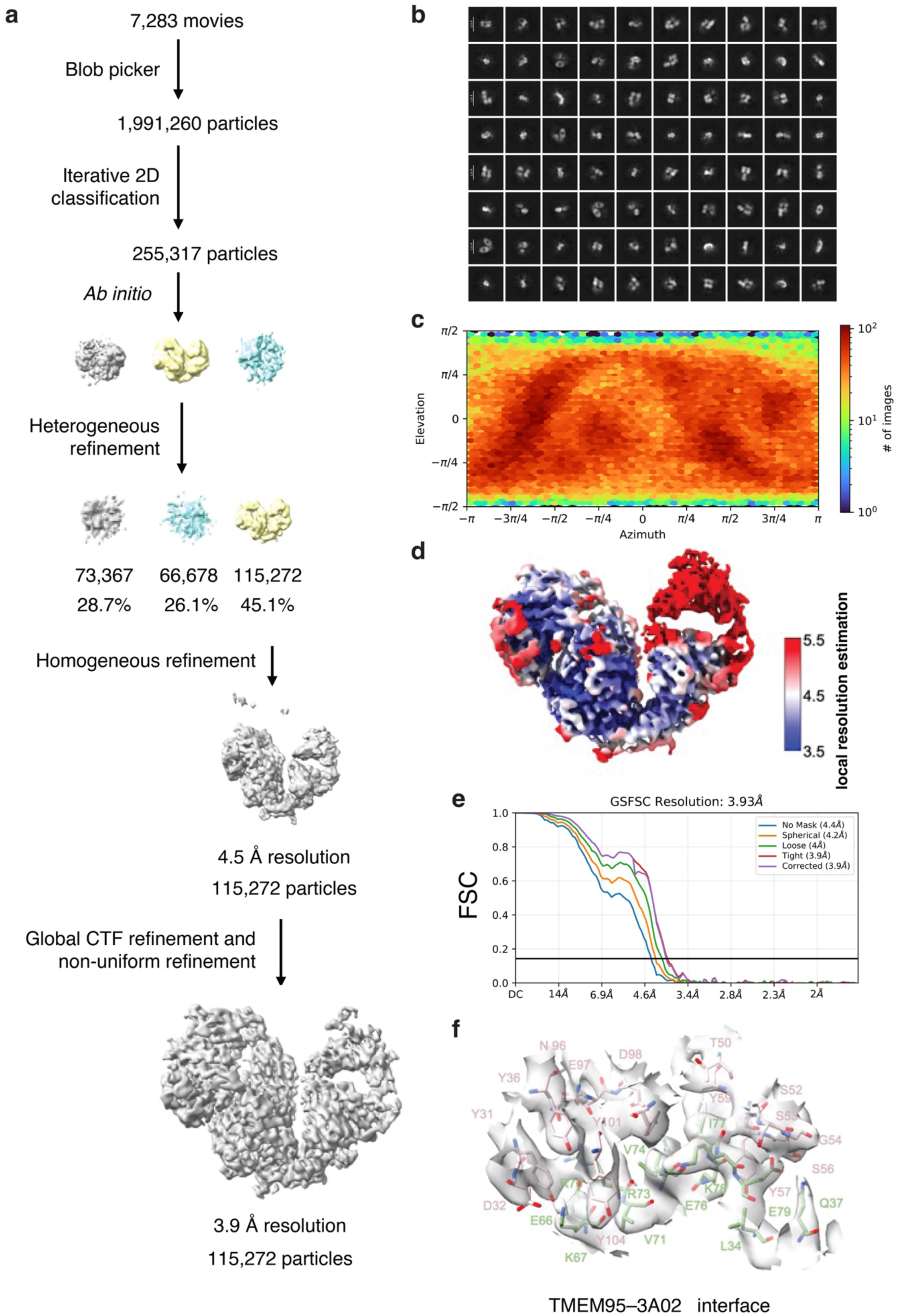
Cryo-EM data processing workflow of the human TMEM95–3A02–6B08 complex. (a) Flow chart of cryo-EM data processing. (b) Representative 2D class averages. Scale bar, 110 Å. (c) Eulerian angle distribution of the particles used in the final 3D reconstruction. (d) Local resolution estimation of the density map. (e) Gold-standard Fourier shell correlation (FSC) curve with the estimated resolution for the overall density map. (f) Local cryo-EM density of the TMEM95–3A02 interface. TMEM95 residues are shown in green; 3A02 residues in pink.

**Extended Data Fig. 7.**
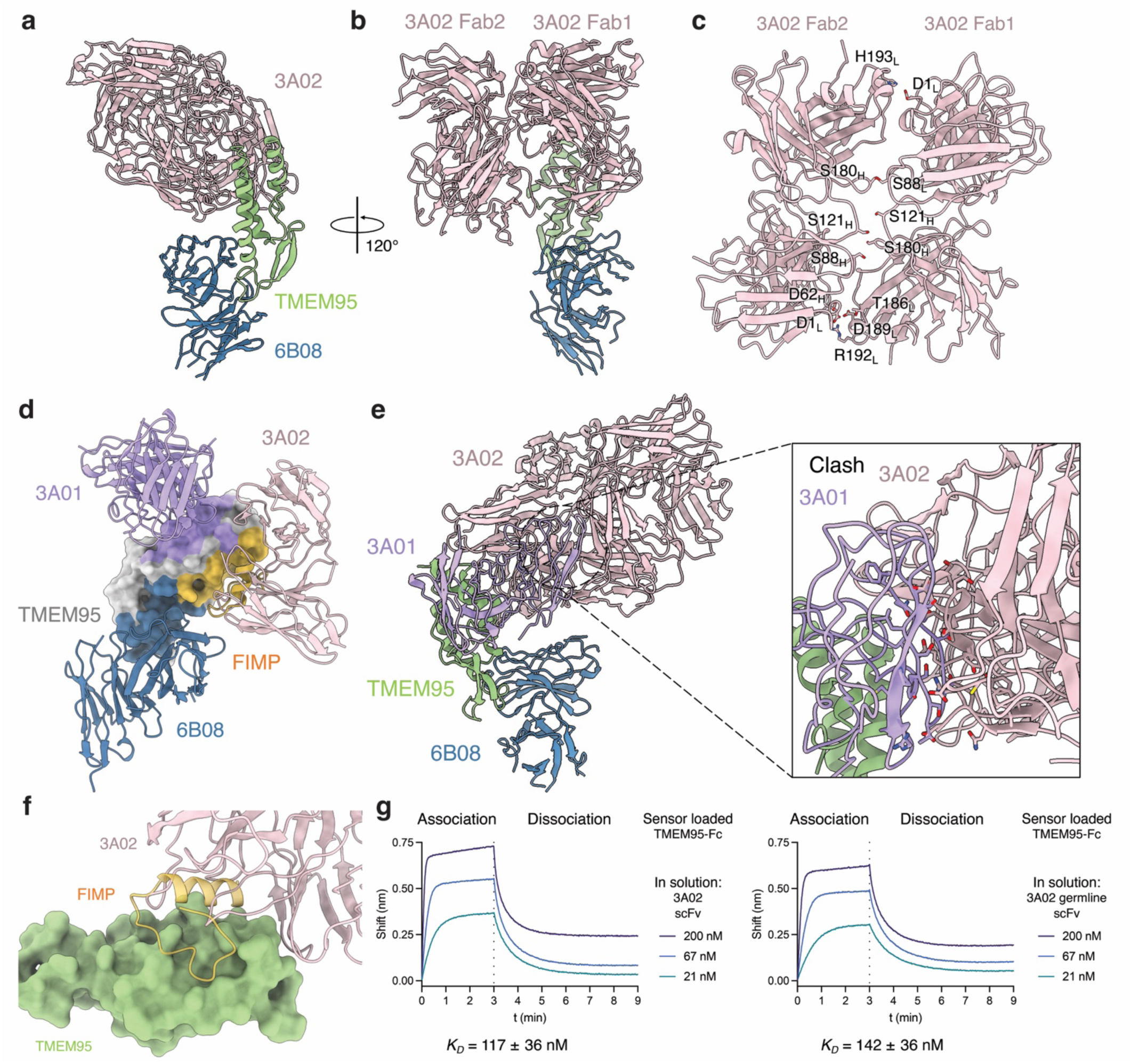
Cryo-EM structure of the human TMEM95–3A02–6B08 complex. (a) Ribbon diagram of the human TMEM95–3A02–6B08 complex. TMEM95 in green, 6B08 in dark blue, and 3A02 in light pink. (b) View of (a) rotated by 120°, with the two 3A02 Fab molecules labeled. (c) Side-chain interactions within a 3A02 Fab dimer. Key interacting residues are shown as sticks. (d) Epitope footprints on TMEM95 of FIMP, 3A01, 6B08, and 3A02. FIMP in yellow, 3A01 in purple, 6B08 in blue, and 3A02 in pink. (e) Superposition of human TMEM95–FIMP–3A01–6B08 and human TMEM95–3A02–6B08 shows that 3A01 sterically clashes with 3A02 Fab2. (f) Superposition of the TMEM95–3A02 and TMEM95–FIMP structures, showing that 3A02 and FIMP occupy overlapping regions on TMEM95. (g) Biolayer interferometry traces showing binding of sensor-loaded TMEM95-Fc to 200 nM, 67 nM, or 21 nM 3A02 scFv or 3A02’s inferred germline scFv. Association, 3 min; dissociation, 6 min.

**Extended Data Fig. 8.**
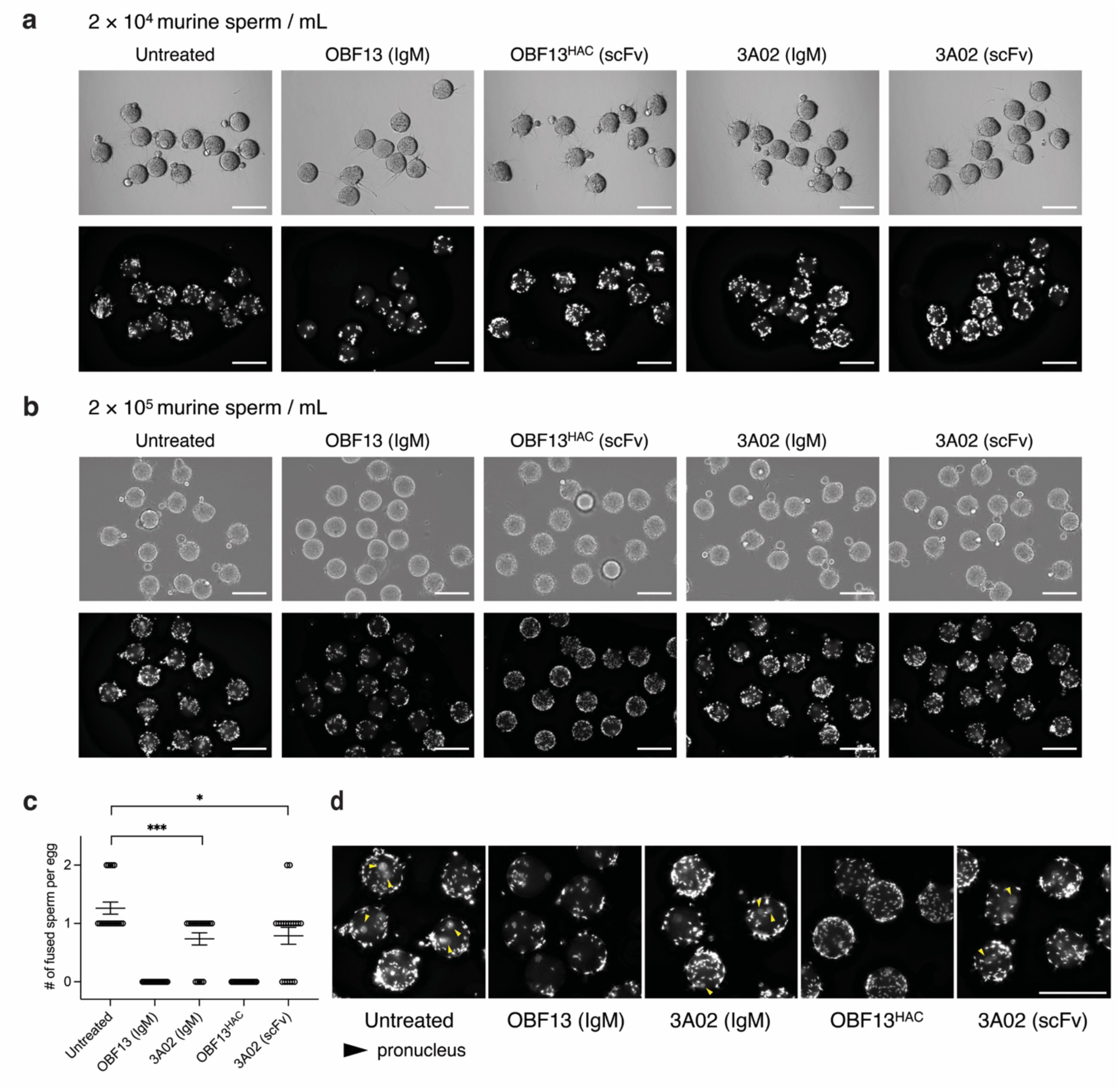
The anti-TMEM95 antibody 3A02 inhibits murine sperm–egg fusion *in vitro*. (a) Representative micrographs of zona-free eggs 6 hours after insemination of 2×10^4^ sperm per mL. DNA was labeled with Hoechst 33342. Sperm were untreated or pretreated with 50 μg/mL OBF13 (IgM), 50 μg/mL OBF13^HAC^ (scFv), 440 μg/mL 3A02 (IgM), or 50 μg/mL 3A02 (scFv). The same set of zoom-in images was shown in Fig. 5b. (b) Representative micrographs of zona-free eggs 6 hours after insemination of 2×10^5^ sperm per mL. (c) Number of fused sperm per zona-free egg (mean ± standard error of the mean, SEM): untreated, 1.3 ± 0.10 (*N* = 19); OBF13 (IgM)-treated, 0 ± 0 (*N* = 19); OBF13^HAC^ (scFv)-treated, 0 ± 0 (*N* = 19); 3A02 (IgM)-treated, 0.74 ± 0.10 (*N* = 19); 3A02 (scFv)-treated, 0.79 ± 0.14 (*N* = 19). (d) The same set of zoom-in images showing the formation of pronuclei inside fertilized eggs. Scale bars, 100 μm.

**Extended Data Fig. 9.**
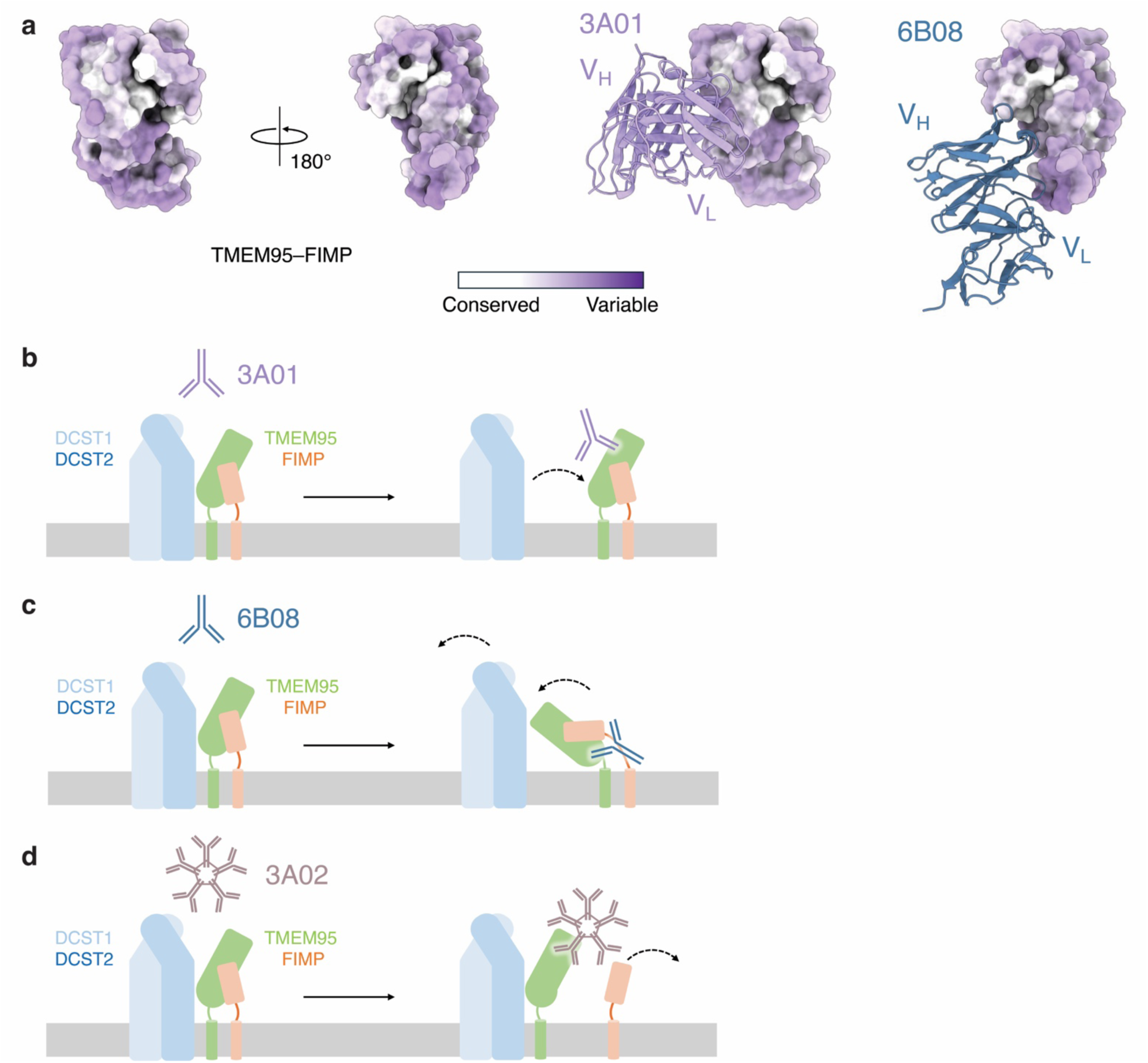
Models of antibody-mediated inhibition of TMEM95. (a) Space-filling models showing sequence conservation on the surface of the TMEM95–FIMP complex, with ribbon diagrams of 3A01 and 6B08 indicating two conserved surfaces outside their epitopes. Cartoon schematics of 3A01-, 6B08-, and 3A02-mediated disruption of the TMEM95–FIMP complex. (b) 3A01 binds the surface of TMEM95 away from the FIMP-binding site, potentially disrupting the interaction of TMEM95–FIMP with the DCST1–DCST2 complex. (c) 6B08 binds the membrane-proximal region of TMEM95, potentially disrupting the orientation of TMEM95–FIMP on the sperm membrane. (d) 3A02 binds the FIMP-binding site of TMEM95, disrupting the TMEM95–FIMP complex.

**Extended Data Fig. 10.**
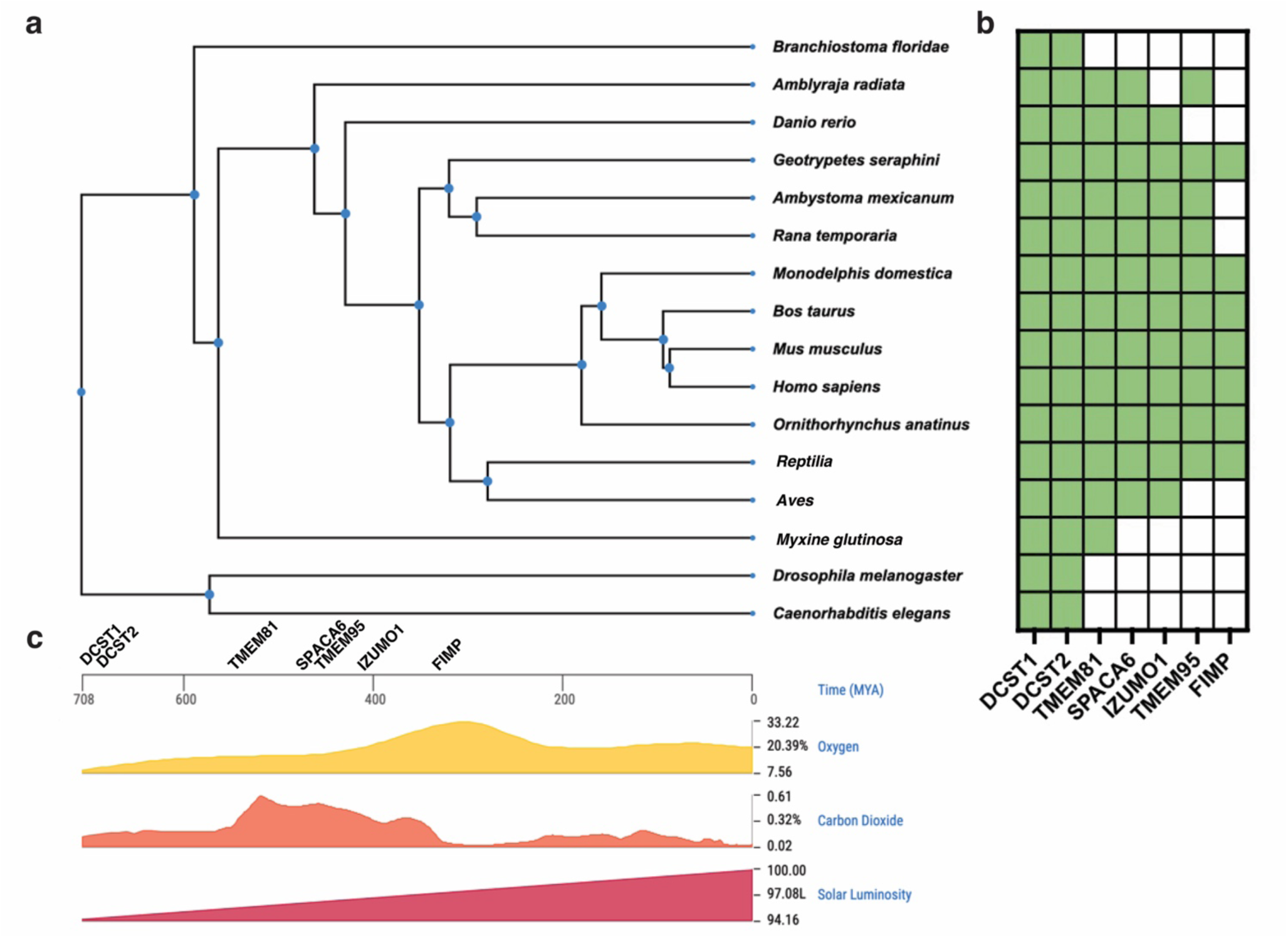
Phylogenetic analysis of gamete fusion-related proteins across the tree of life. (a) PSI-BLAST searches of the NCBI database using IZUMO1, SPACA6, TMEM81, TMEM95, FIMP, DCST1, and DCST2 as queries were used to determine where in the tree of life these sequences are found. Phylogenetic relationships of representative species were generated using the TimeTree database. TMEM95 appears in vertebrates before FIMP. (b) The potential emergence of sperm fertilization-essential proteins is plotted along a geological time axis. (c) Changes in atmospheric oxygen levels, carbon dioxide concentrations, and light availability across evolutionary time.

**Extended Data Table 1.** Plasmids and yeast strains used in this study.

| **Plasmids for human Expi293F cell line protein expression** |  |  |
| --- | --- | --- |
| **Plasmid** | **Description** | **Reference** |
| pPL85 | pADD2 hFIMP_F93A-ΔN-α-helix-Fc Holes | This study |
| pPL92 | pADD2 hTMEM95-α-helix-Fc-His<sub>6</sub> Knobs | This study |
| pPL97 | pVRC TMEM95 3A02 Heavy Fab | This study |
| pPL98 | pVRC TMEM95 3A02 Light Fab | This study |
| pPL112 | pADD2 3A02 scFv-His<sub>6</sub> | This study |
| pPL117 | pADD2 3A02 germline scFv-His<sub>6</sub> | This study |
| pST1363 | pADD2 hFIMP-Fc-Avi-His<sub>6</sub> | This study |
| pST1359 | pADD2 hTMEM95-Fc-Avi-His<sub>6</sub> | (1) |
| pST1365 | pADD2 mFIMP-Fc-Avi-His<sub>6</sub> | This study |
| pST1557 | pADD2 hTMEM95-Ctag | (1) |
| pST1679 | pCAG mTMEM95-3FLAG-TM | This study |
| pST1720 | pVRC hTMEM95 3A01 mIgG1 Heavy | (1) |
| pST1721 | pVRC hTMEM95 3A01 mIgG1 Fab Heavy | (1) |
| pST1722 | pVRC hTMEM95 3A01 Light | (1) |
| pST1723 | pVRC hTMEM95 6B08 mIgG1 Heavy | (1) |
| pST1724 | pVRC hTMEM95 6B08 mIgG1 Fab Heavy | (1) |
| pST1725 | pVRC hTMEM95 6B08 Light | (1) |
| pST1735 | pADD2 OBF13_HAC scFv-His<sub>6</sub> | (2) |
| pST1763 | pADD2 hTMEM95_R110A-Fc-Avi-His<sub>6</sub> | This study |
| pST2341 | pADD2 mTMEM95-Fc-Avi-His<sub>6</sub> Knobs | This study |
| pST2344 | pADD2 mFIMP-Fc-Avi-His<sub>6</sub> Holes | This study |
| pST2359 | pADD2 hFIMP_F66A-Fc-Avi-His<sub>6</sub> | This study |
| pST2360 | pADD2 hFIMP_F69A-Fc-Avi-His<sub>6</sub> | This study |
| pST2361 | pADD2 hFIMP_Y71A-Fc-Avi-His<sub>6</sub> | This study |
| pST2362 | pADD2 hFIMP_D73A-Fc-Avi-His<sub>6</sub> | This study |
| pST2363 | pADD2 hFIMP_F85A-Fc-Avi-His<sub>6</sub> | This study |
| pST2364 | pADD2 hFIMP_I86A-Fc-Avi-His<sub>6</sub> | This study |
| pST2365 | pADD2 hFIMP_E88A-Fc-Avi-His<sub>6</sub> | This study |
| pST2366 | pADD2 hFIMP_F93A-Fc-Avi-His<sub>6</sub> | This study |

“h” is for human and “m” is for mouse

**Extended Data Table 2.**
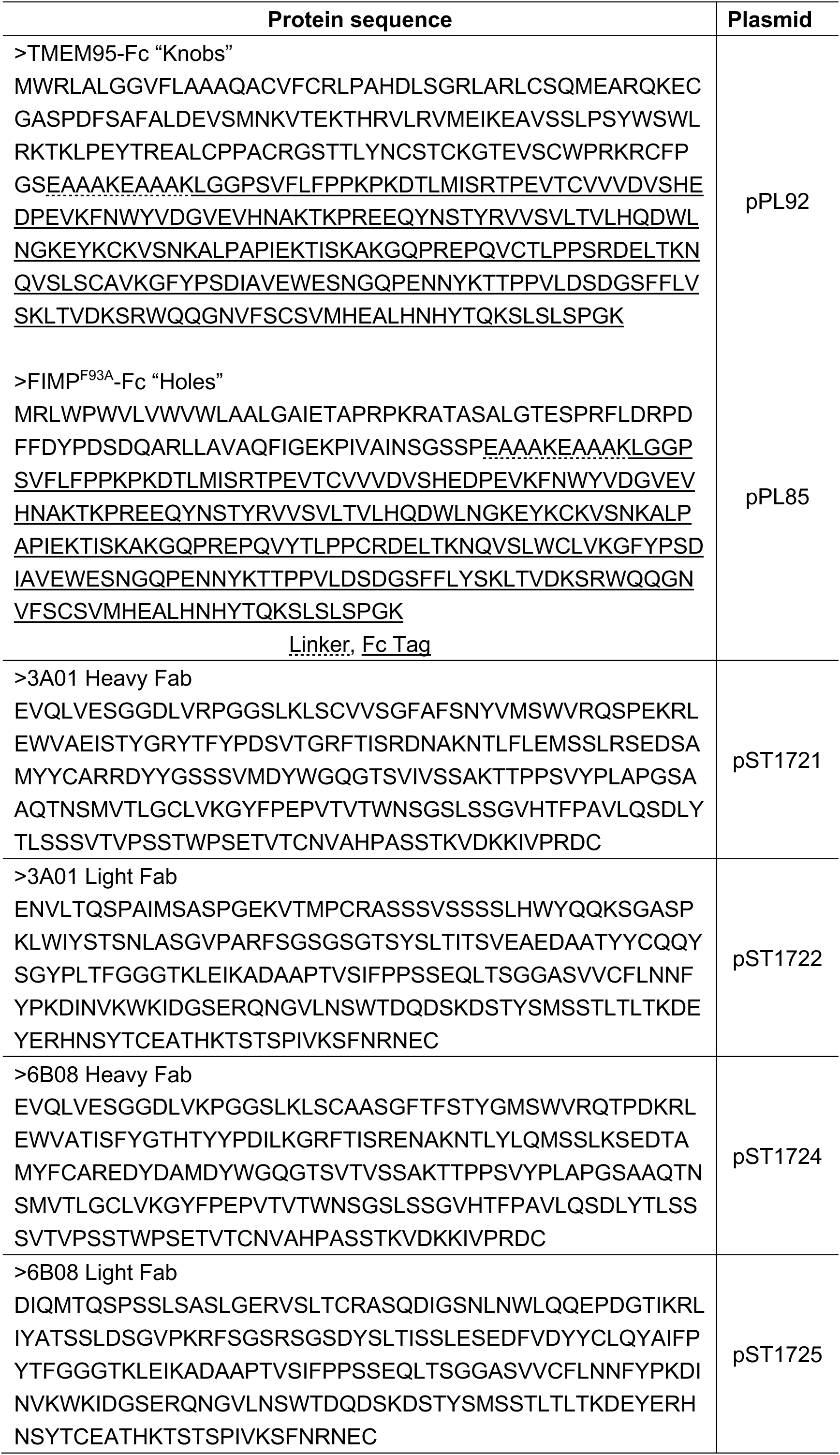
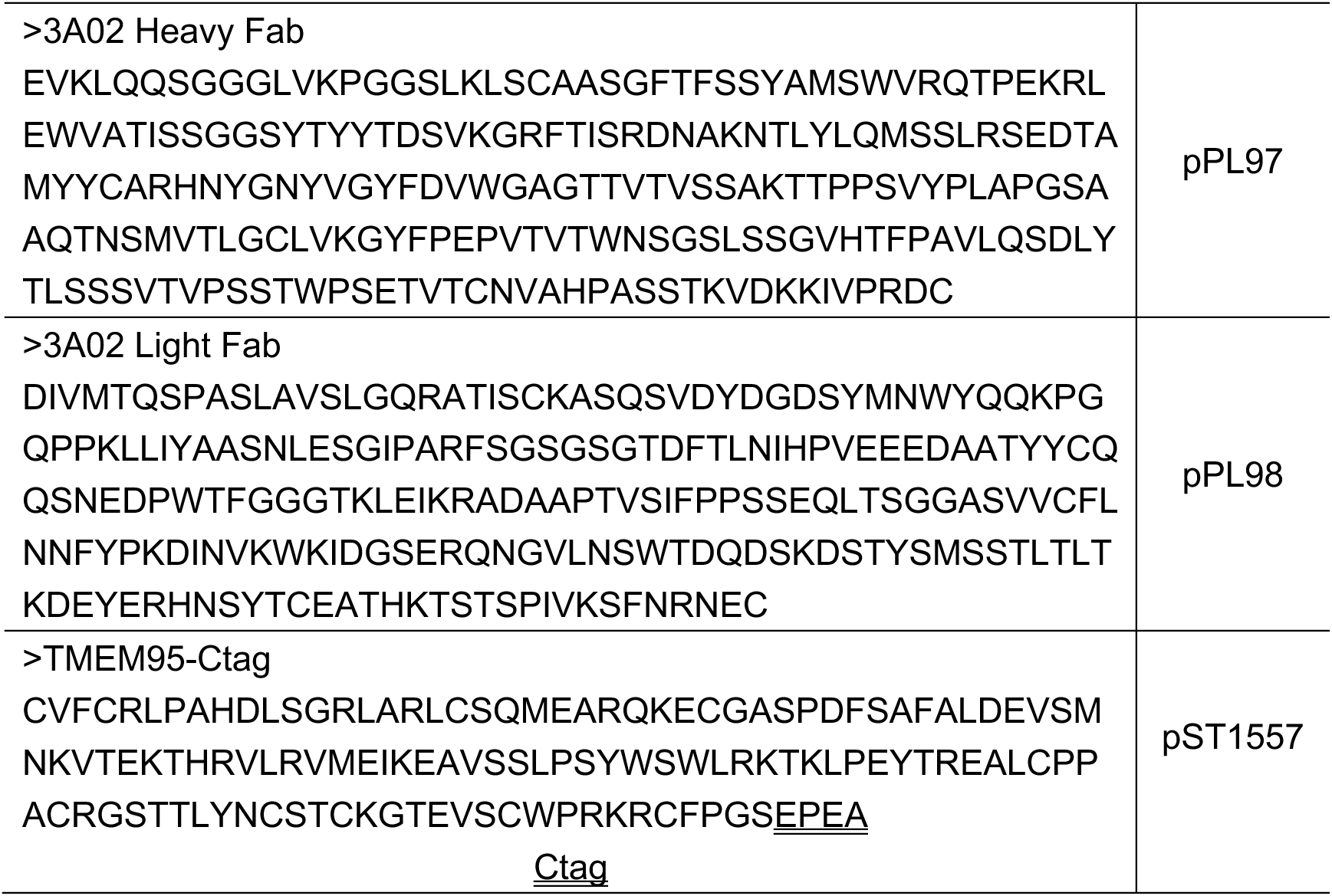
Protein sequences used for cryo-EM in this study.

## References

1 Lorenzetti, D. et al. A transgenic insertion on mouse chromosome 17 inactivates a novel immunoglobulin superfamily gene potentially involved in sperm-egg fusion. Mamm. Genome 25, 141–148, doi:10.1007/s00335-013-9491-x (2014).

2 Noda, T. et al. Sperm proteins SOF1, TMEM95, and SPACA6 are required for sperm-oocyte fusion in mice. Proceedings of the National Academy of Sciences of the United States of America, doi:10.1073/pnas.1922650117 (2020).

3 Lamas-Toranzo, I. et al. TMEM95 is a sperm membrane protein essential for mammalian fertilization. eLife 9, doi:10.7554/eLife.53913 (2020).

4 Barbaux, S. et al. Sperm SPACA6 protein is required for mammalian Sperm-Egg Adhesion/Fusion. Sci Rep 10, 5335, doi:10.1038/s41598-020-62091-y (2020).

5 Noda, T. et al. Sperm membrane proteins DCST1 and DCST2 are required for sperm-egg interaction in mice and fish. Commun Biol 5, 332, doi:10.1038/s42003-022-03289-w (2022).

6 Pausch, H. et al. A nonsense mutation in TMEM95 encoding a nondescript transmembrane protein causes idiopathic male subfertility in cattle. PLoS Genet. 10, e1004044, doi:10.1371/journal.pgen.1004044 (2014).

7 Tang, S. et al. Human sperm TMEM95 binds eggs and facilitates membrane fusion. Proceedings of the National Academy of Sciences of the United States of America 119, e2207805119, doi:10.1073/pnas.2207805119 (2022).

8 Inoue, N., Hagihara, Y. & Wada, I. Evolutionarily conserved sperm factors, DCST1 and DCST2, are required for gamete fusion. eLife 10, doi:10.7554/eLife.66313 (2021).

9 Deneke, V. E. et al. A conserved fertilization complex bridges sperm and egg in vertebrates. Cell, doi:10.1016/j.cell.2024.09.035 (2024).

10 Inoue, N., Ikawa, M., Isotani, A. & Okabe, M. The immunoglobulin superfamily protein Izumo is required for sperm to fuse with eggs. Nature 434, 234–238, doi:10.1038/nature03362 (2005).

11 Inoue, N. & Wada, I. Deletion of the initial methionine codon of the Tmem95 gene causes subfertility, but not complete infertility, in male mice. Biol. Reprod. 106, 378–381, doi:10.1093/biolre/ioab246 (2022).

12 Chang, H. Y., Gierke, T., Tang, S. & Lu, Y. Molecular interplay between sperm and oocyte: a narrative review. Hum. Reprod. Update, doi:10.1093/humupd/dmag008 (2026).

13 Bianchi, E., Doe, B., Goulding, D. & Wright, G. J. Juno is the egg Izumo receptor and is essential for mammalian fertilization. Nature 508, 483–487, doi:10.1038/nature13203 (2014).

14 Elofsson, A., Han, L., Bianchi, E., Wright, G. J. & Jovine, L. Deep learning insights into the architecture of the mammalian egg-sperm fusion synapse. eLife 13, doi:10.7554/eLife.93131 (2024).

15 Vance, T. D. R. et al. SPACA6 ectodomain structure reveals a conserved superfamily of gamete fusion-associated proteins. Commun Biol 5, 984, doi:10.1038/s42003-022-03883-y (2022).

16 L, G.-B., Jg, H., I, L.-T., M, J.-M. & P, B.-A. The Sperm Olfactory Receptor OLFR601 is Dispensable for Mouse Fertilization. Front Cell Dev Biol 10, 854115, doi:10.3389/fcell.2022.854115 (2022).

17 Fujihara, Y. et al. Spermatozoa lacking Fertilization Influencing Membrane Protein (FIMP) fail to fuse with oocytes in mice. Proceedings of the National Academy of Sciences of the United States of America 117, 9393–9400, doi:10.1073/pnas.1917060117 (2020).

18 V. E. Deneke et al. The SPARK complex forms the molecular basis of vertebrate fertilization. bioRxiv, doi:doi:10.64898/2026.05.14.724128 (2026).

19 J. N. Elango, I. Shin, A. Gurjar & Krauchunas, A. R. Characterization of spe-40/Fam187 identifies a deeply conserved sperm protein at the C. elegans fertilization synapse. bioRxiv, doi:10.64898/2026.05.14.723898 (2026).

20 Lu, Y. et al. 1700029I15Rik orchestrates the biosynthesis of acrosomal membrane proteins required for sperm-egg interaction. Proceedings of the National Academy of Sciences of the United States of America 120, e2207263120, doi:10.1073/pnas.2207263120 (2023).

21 Lu, Y., Ikawa, M. & Tang, S. Allosteric inhibition of the IZUMO1-JUNO fertilization complex by the naturally occurring antisperm antibody OBF13. Proceedings of the National Academy of Sciences of the United States of America 122, e2425952122, doi:10.1073/pnas.2425952122 (2025).

22 Hie, B. L. et al. Efficient evolution of human antibodies from general protein language models. Nat. Biotechnol. 42, 275–283, doi:10.1038/s41587-023-01763-2 (2024).

23 Dal Porto, J. M., Haberman, A. M., Kelsoe, G. & Shlomchik, M. J. Very low affinity B cells form germinal centers, become memory B cells, and participate in secondary immune responses when higher affinity competition is reduced. J. Exp. Med. 195, 1215–1221, doi:10.1084/jem.20011550 (2002).

24 Shih, T. A., Meffre, E., Roederer, M. & Nussenzweig, M. C. Role of BCR affinity in T cell dependent antibody responses in vivo. Nat. Immunol. 3, 570–575, doi:10.1038/ni803 (2002).

25 Shatz, W. et al. Knobs-into-holes antibody production in mammalian cell lines reveals that asymmetric afucosylation is sufficient for full antibody-dependent cellular cytotoxicity. MAbs 5, 872–881, doi:10.4161/mabs.26307 (2013).

26 Wang, Z. et al. Universal PCR amplification of mouse immunoglobulin gene variable regions: the design of degenerate primers and an assessment of the effect of DNA polymerase 3′ to 5′ exonuclease activity. Journal of Immunological Methods 233, 167–177, 10.1016/S0022-1759(99)00184-2 (2000).

27 Ashkenazy, H. et al. ConSurf 2016: an improved methodology to estimate and visualize evolutionary conservation in macromolecules. Nucleic acids research 44, W344–350, doi:10.1093/nar/gkw408 (2016).

1. S. Tang et al., Human sperm TMEM95 binds eggs and facilitates membrane fusion. Proceedings of the National Academy of Sciences 119, e2207805119 (2022).

2. Y. Lu, M. Ikawa, S. Tang, Allosteric inhibition of the IZUMO1–JUNO fertilization complex by the naturally occurring antisperm antibody OBF13. Proceedings of the National Academy of Sciences 122, e2425952122 (2025).

